# Locus-resolved expression and manipulation of human-specific transposable elements reveal their role in timely progression of early brain development

**DOI:** 10.64898/2026.11.20.768047

**Authors:** Laura Deelen, Sophie A Marlow, Paulina Urbanaviciute, Tabitha T Richards, Zoe H Mitchell, Martina Demurtas, Andria Koulle, Xiao Li, Kathryn R Bowles, Rebecca V Berrens, Marco Trizzino

## Abstract

The human cerebral cortex has undergone remarkable expansion during primate evolution, yet the regulatory innovations underlying human-specific neurodevelopmental programmes remain poorly understood. Transposable elements (TEs) comprise approximately half of the human genome and represent a major source of lineage-specific regulatory novelty, but their individual contributions to cortical development have been inaccessible due to the technical challenge of resolving near-identical repetitive sequences. Here we apply single-cell long-read sequencing (CELLO-seq) to generate the first locus-resolved atlas of young human-specific TE expression in cortical organoids, revealing that individual loci from multiple TE families, including SVAs, LTR5Hs, and L1Hs, are expressed across all major cortical cell types and carry binding motifs for key cortical and neurogenic transcription factors. Using CRISPRi to silence SVAs and LTR5Hs during organoid differentiation, we demonstrate that their combined loss causes coordinated failure of the radial glia-to-neuron transition, characterised by progenitor retention, transcriptional ventralisation, impaired synaptic maturation and a differentiation block in interneuron precursors. Co-expression network and regulon analyses identify loss of dorsal cortical identity programmes and gain of ventralisation-associated transcriptional activity as the dominant molecular consequences of TE silencing. Motif-to-regulon convergence directly links SVA and LTR5Hs chromatin accessibility to the loss of specific TF binding programmes. These findings establish young and human-specific TEs as functional components of the regulatory architecture governing the timing of cortical neurogenesis, with implications for understanding human brain evolution and neurodevelopmental disease.

## Introduction

The human cerebral cortex is distinguished from that of other primates by its dramatic expansion in size and complexity, a transformation that unfolded over a relatively short evolutionary timescale and is thought to reflect, at least in part, the emergence of novel gene regulatory programmes unique to the human lineage. Transposable elements (TEs), which comprise approximately half of the human genome, have increasingly been recognised as a major source of such regulatory novelty^1–7^. Through their capacity to insert into new genomic locations, TEs can shape new transcription factor binding sites and introduce novel promoters and enhancers, reshaping the regulatory landscape of their host genomes in ways that may alter gene expression in a lineage-specific manner^8–11^. Although most TEs are epigenetically silenced, a subset remains accessible and transcriptionally active, particularly during development, when chromatin is broadly permissive and cell fate decisions are being made. In primates, evolutionarily young TEs have been shown to rewire gene regulatory networks during embryogenesis, placentation, and neural development, making them compelling candidates for contributing to the regulatory innovations that underlie human-specific brain development^9–14^.

Among the youngest and most human-relevant TE families are the SINE-Vntr-Alus (SVAs), LTR5Hs, and LINE-1 subfamilies L1Hs and L1PA2/3, all of which contain members that arose during or after the human-chimpanzee split approximately six million years ago^15–19^. SVAs in particular represent the most recently active TE family in the human genome and have been implicated as cis-regulatory elements in hippocampal neurogenesis, neural crest development, and pluripotent stem cells^12–14,20^. LTR5Hs share an ancestral origin with SVAs through the endogenous retrovirus HERVK-10, making them a closely related retroviral TE family whose regulatory potential during human neurodevelopment has remained largely unexplored alongside SVAs. Despite growing evidence for their regulatory importance, human-specific TEs remain poorly characterised because their recent evolutionary origin means many copies retain near-identical sequences, rendering them inaccessible to conventional short-read sequencing, which cannot distinguish between individual loci^21^. As a result, most studies average signals across entire TE subfamilies, masking the distinct genomic contexts and regulatory functions of individual insertions.

Here we address this longstanding limitation in the field by applying single CELl LOng-read sequencing (CELLO-seq), a single-cell long-read sequencing approach that captures full-length transcripts and enables locus-resolved quantification of TE expression^22,23^, to cortical organoids modelling early human forebrain development. We analysed cortical organoids at day-21 of differentiation, a stage corresponding to active neurogenesis and the transition from radial glia to early neurons. The human cerebral cortex has undergone substantial expansion compared to other primates, and evolutionarily young TEs may have contributed to the regulatory programs underlying human-specific neurodevelopment^24–26^.

With this approach, we generated the first locus-resolved atlas of young and human-specific TE expression in cortical organoids and identified the transcription factor binding repertoire of expressed SVAs and LTR5Hs. To directly test the functional requirement of these elements for cortical neurogenesis, we employed a previously published inducible CRISPR (CRISPRi) system to silence SVAs and LTR5Hs^13,14,20^ during cortical organoid differentiation. We characterised the transcriptional consequences at single-cell resolution using differential expression analysis, pseudotime and RNA velocity trajectory modelling, and high-dimensional weighted gene co-expression network analysis. Together, our findings reveal that human-specific TEs function as enhancers occupied by key dorsal cortical identity transcription factors during early neurogenesis. Their silencing causes a coordinated failure of the radial glia-to-neuron transition, characterised by progenitor retention, transcriptional ventralisation, and a differentiation block. Collectively, this implicates SVAs and LTR5Hs as functional components of a human-specific regulatory architecture that governs the timing of cortical neurogenesis.

## Results

### CELLO-seq enables locus-level detection of young transposable element expression

To characterise evolutionarily young TEs active during early human cortical development, we employed a recently optimised cortical organoid model^27^. This model directs human induced pluripotent stem cells (hiPSCs) through sequential neuronal induction (d0-5), proliferation (d6-24) and maturation media (>d25) without the need for an embedding matrix. In addition, the use of slit-well plates, which allows shared media between all 96 wells but ensures physical separation, warrants increased reproducibility (**Fig. 1a,b**). This system produced cortical organoids of consistent size and growth trajectory for the two lines used in this study (-gRNAs/+gRNAs, see Methods) (**Fig. 1c,d**). At d21, we detected gene expression of key cell type markers including neural progenitor marker *PAX6*, dorsal forebrain marker *FOXG1*, deep layer neuron markers *TBR1* and *CTIP2/BCL11B*, pan-neuronal marker *MAP2*, and negative control *COL1A2* (**Fig. 1e**). Protein levels for PAX6, SOX2, FOXG1, TBR1, CTIP2 and TUJ1 were confirmed at both d21 and 4 months (**Fig. 1f**).

**Figure 1.**
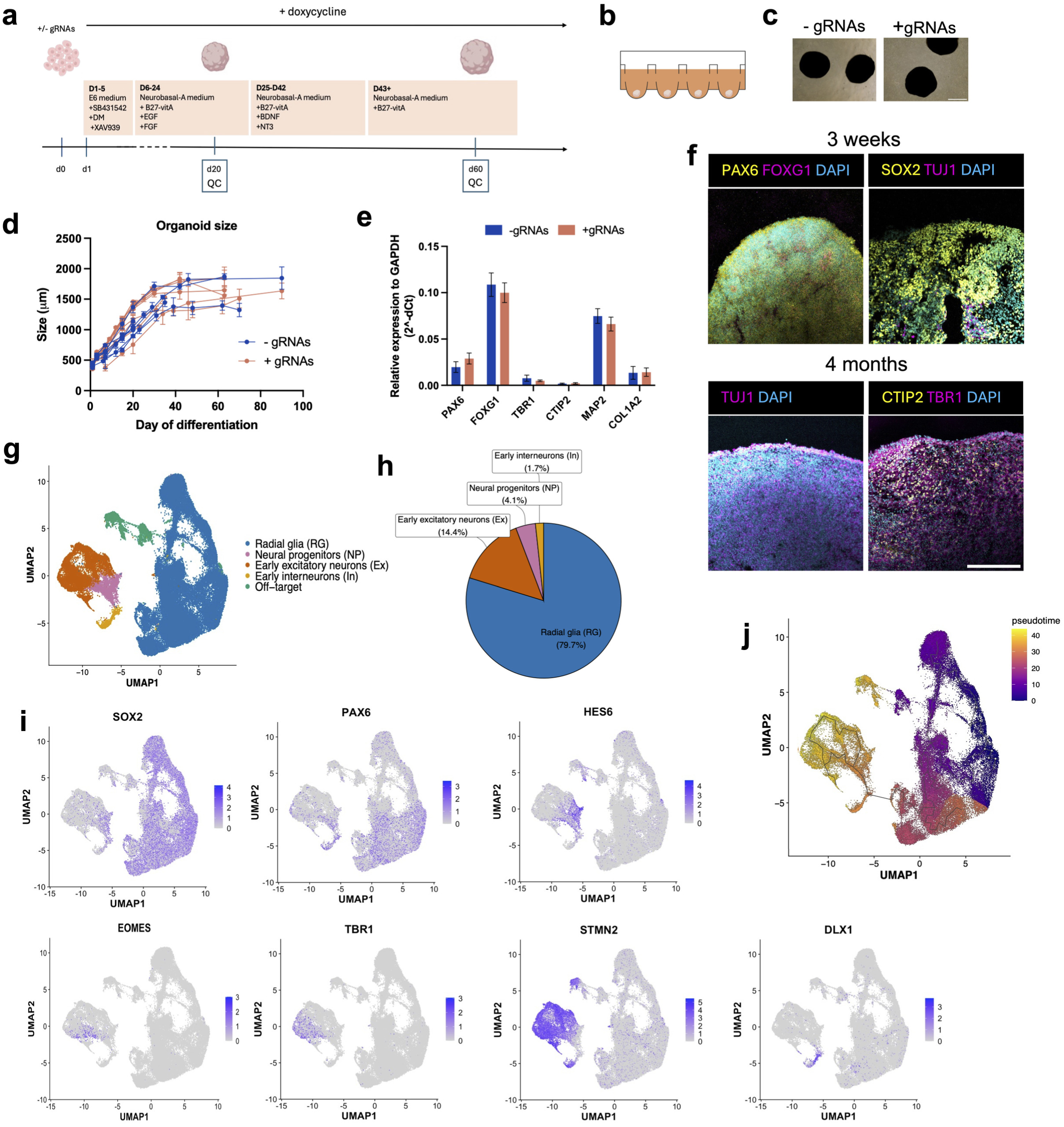
Cortical organoid model and single-cell transcriptomic characterisation. **(a)** Schematic of the cortical organoid differentiation protocol. Human iPSCs engineered with an inducible dCas9-KRAB system, with or without gRNAs targeting SVAs and LTR5Hs (-/+ gRNAs) are differentiated according to Bertucci et al., 2022. Doxycycline is added from d1 to induce dCas9-KRAB expression throughout differentiation. Quality control (QC) is performed around d20 and d60. **(b)** Schematic of slit-well plates used for organoid culture, enabling shared media between wells and increased reproducibility, **(c)** Representative images of cortical organoids at three weeks, showing comparable morphology between gRNAs and ÷gRNAs lines. Scale bar: 1000 pm. (d) Organoid size (pm) measurements showing consistent growth trajectories between conditions. Error bars indicate standard deviation, **(e)** Quantitative analysis shows no significant differences in gene expression of dorsal forebrain markers PAX6 and FOXG1, cortical layers VI (TBR1) and V (CTIP2), the neuronal marker MAP2 or the negative control COL1A2. n=16-19. (f) Schematic of marker visualisation. Confirmation of dorsal forebrain markers PAX6, FOXG1, SOX2, and low levels of early neuron marker TUJ1 at d20. At four months, sections were positive for deep-layer markers TBR1, CTIP2 and higher levels of TUJ1. Scale bar, 200 pm. (g) UMAP visualisation of single-cell RNA sequencing data from d21 cortical organoids, **(h)** Pie chart showing the proportional composition of cell types at d21. **(i)** Feature plots showing the expression of canonical cell type marker genes on the UMAP, including radial glia (RG) markers SOX2, PAX6, neuronal progenitor (NP) markers HES6 and EOMES, deep-layer marker TBR1, pan-neuronal marker STMN2, interneuron marker DLX1, and the off-target population markers SIX1 (placodes) and SOX10 (neural crest), (j) UMAP visualisation coloured by Monocle3 pseudotime values, confirming the expected differentiation trajectory from radial glia (early pseudotime, dark purple) through neural progenitors to excitatory neurons (late pseudotime, yellow).

To capture early neurogenesis, we focused on d21 organoids and characterised their cellular composition by single cell RNA sequencing (scRNA-seq) on two independent batches of five pooled organoids per cell line. Following data normalisation, dimensionality reduction and clustering, cell type annotation was assigned using scRNA-seq reference expression data from the human neocortex at gestational week (GW) 17-18^28^, consistent with previous use of this protocol^27,29^, and GW6-10^30^. At d21, the expected majority of cells were radial glia (RG), with neuronal progenitors (NP), early excitatory neurons (Ex) and early interneurons (In) also present, reflecting the predominantly neurogenic stage of these organoids (**Fig. 1g-i**). As a common observation in cortical organoids, we also observed a minor amount of off-target neural crest (*SOX10*) and placodes (*SIX1*) populations, which were excluded from further analyses (**Supp. Fig.1**). Pseudotime trajectory analysis (Monocle3) confirmed the expected differentiation axis from radial glia through neuronal progenitors to early neurons (**Fig. 1j**).

To detect young TEs expressed during early brain development, we performed CELLO-seq on five pooled cortical organoids at d21 of differentiation. CELLO-seq captures full-length transcripts at single-cell resolution, optimised for repetitive sequences such as TEs and isoform detection^22^. We profiled 525 individual cells using this approach, and despite the inherently lower cell numbers amenable to long-read sequencing, CELLO-seq broadly preserved the UMAP topology and cell type representation observed by short-read scRNA-seq (**Fig. 2a-c**). Critically, compared to our short-read scRNAseq data, we observed a six-to-ten-fold increase in TE detection signal with CELLO-sequencing when pseudobulking counts for young TE elements (**Fig. 2d**). Even for older elements, CELLO-sequencing increased the detection signal 4-fold on average (**Supp. Fig. 2**). This increase reflects the ability of long reads to span entire repetitive elements and uniquely assign transcripts to individual loci, a fundamental limitation of short-read approaches. CELLO-seq detected transcripts from various human-specific TEs of different TE family origin, whose loci were confirmed accessible by bulk ATAC-seq data of d21 organoids (**Fig. 2e**).

**Figure 2.**
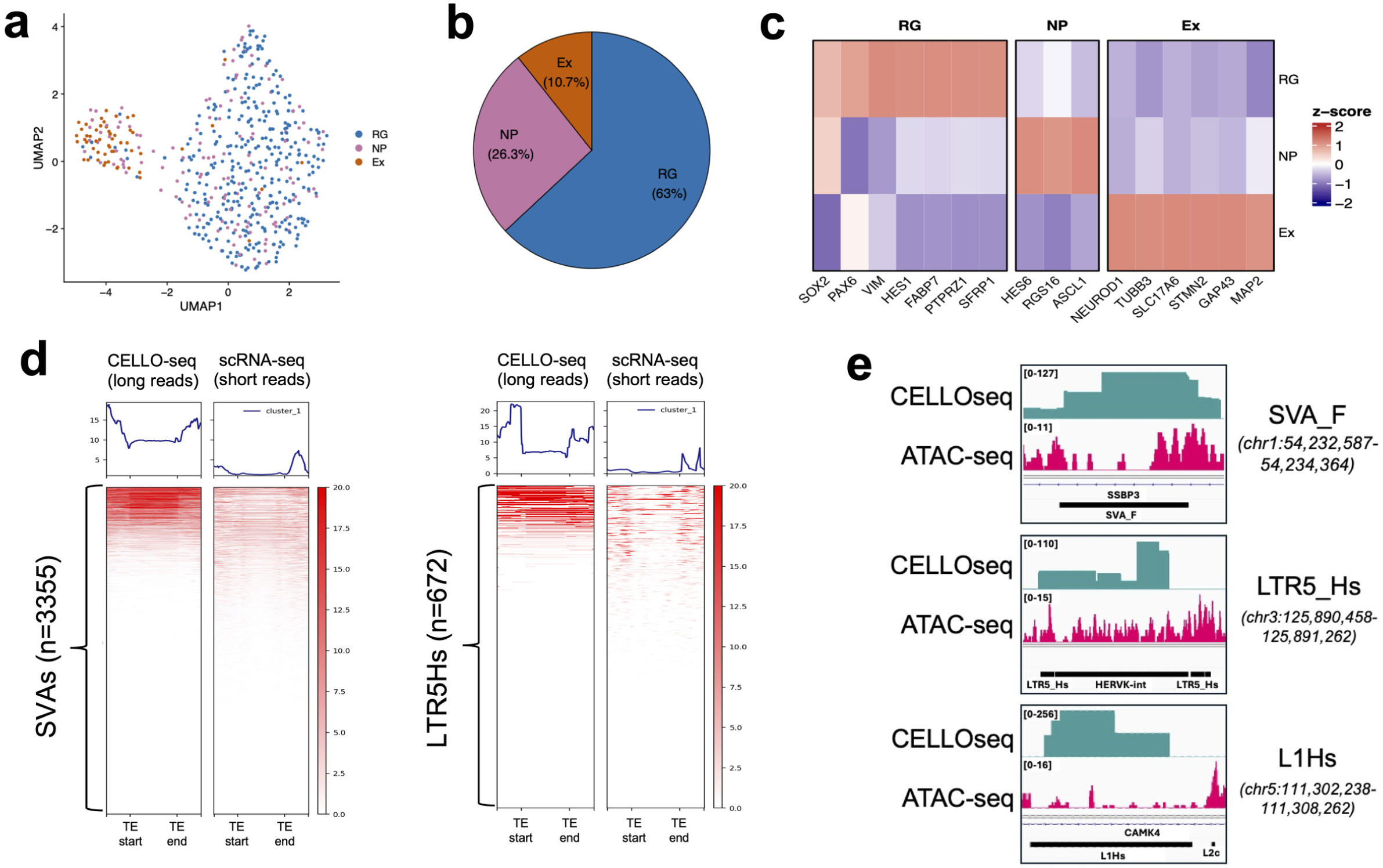
CELLO-seq enables locus-resolved detection of young human-specific transposable element expression in cortical organoids. **(a)** UMAP visualisation of CELLO-seq data from d21 cortical organoids (n=525 cells), demonstrating preservation of the UMAP topology observed by short-read scRNA-seq. **(b)** Cell type composition of the d21 CELLO-seq dataset, **(c)** Heatmap of z-scored average expression of cell type marker genes across RG, NP, and Ex populations in the CELLO-seq dataset, **(d)** Heatmaps of read coverage across SVA loci and LTR5Hs loci reveal six-to-tenfold increase in coverage of young TEs with CELLO-seq compared to short-read sequencing, **(e)** Genome browser tracks showing CELLO-seq expression signal and bulkATAC-seq accessibility signal at three representative human-specific TE loci.

### Human-specific TE expression is enriched in radial glia and distributed genome-wide

To focus on TEs with potential human-specific regulatory functions in the developing brain, we next identified which expressed elements are absent from other great ape genomes (**Fig. 3a**). We restricted our analysis to recently active retrotransposon families known to contain human-specific or evolutionarily young members: SVAs (A-F/F1), LTR5Hs, L1Hs, L1PA2 and L1PA3. For each element within these subfamilies, we identified the 5 kb flanking regions on either side of the element in the human reference genome (hg38), and performed LiftOver to the chimpanzee, gorilla and orangutan genomes to find the corresponding sequences in their respective genomes. If these 5 kb flanking regions were mapped adjacently to each other (gap ≤ 25% of the element length), the TE was defined as absent in that species, and vice versa, if these flanking regions were found at a distance consistent with the TE insertion (≥60% of the element length), the TE was defined as present in that genome (see Methods). All elements were then labelled accordingly to being either human-specific, shared with other great ape species, or ambiguous. Crossing this classification with the CELLO-seq expression data yielded a catalogue of human-specific TEs actively transcribed during early cortical development (labelled in orange) (**Fig. 3b, Supp. File 1**). SVAs represent the youngest TE family in the human genome and consist of seven subfamilies (SVA_A-F/F1), of which the oldest (SVA_A-D) are shared across great apes, having arisen approximately 25 Million years ago (Mya), whereas the youngest subfamilies (SVA_E,F) have been deemed human-specific as these elements arose during or after the human-chimpanzee split (∼6 Mya)^31^. We detected tens of different SVAs per subfamily members expressed in the brain, both from the younger subfamilies SVA_D (116 loci, 7.25%), SVA_E (19 loci, 2.62%) and SVA_F (49 loci, 4.65%), as well as the older subfamilies with predominantly gorilla/orangutan-shared elements, SVA_A (20 loci, 1.71%), SVA_B (32 loci, 3.62%) and SVA_C (26 loci, 4.85%) (**Fig. 3b**). To our knowledge, this represents the first systematic, locus-level quantification of young TE expression in cortical organoids. We detect endogenous SVA expression across all seven subfamilies, providing a reference landscape of locus-specific SVA transcriptional activity during early human neurogenesis that has previously been unobtainable. Among other TE subfamilies harbouring human-specific elements, expressed loci were likewise identified: LTR5Hs (29 loci, 4.04%) and L1Hs (160 loci, 9.28%), as well as the older L1PA2 (275 loci, 5.24%) and L1PA3 (408 loci, 3.58%) (**Fig. 3b**).

**Figure 3.**
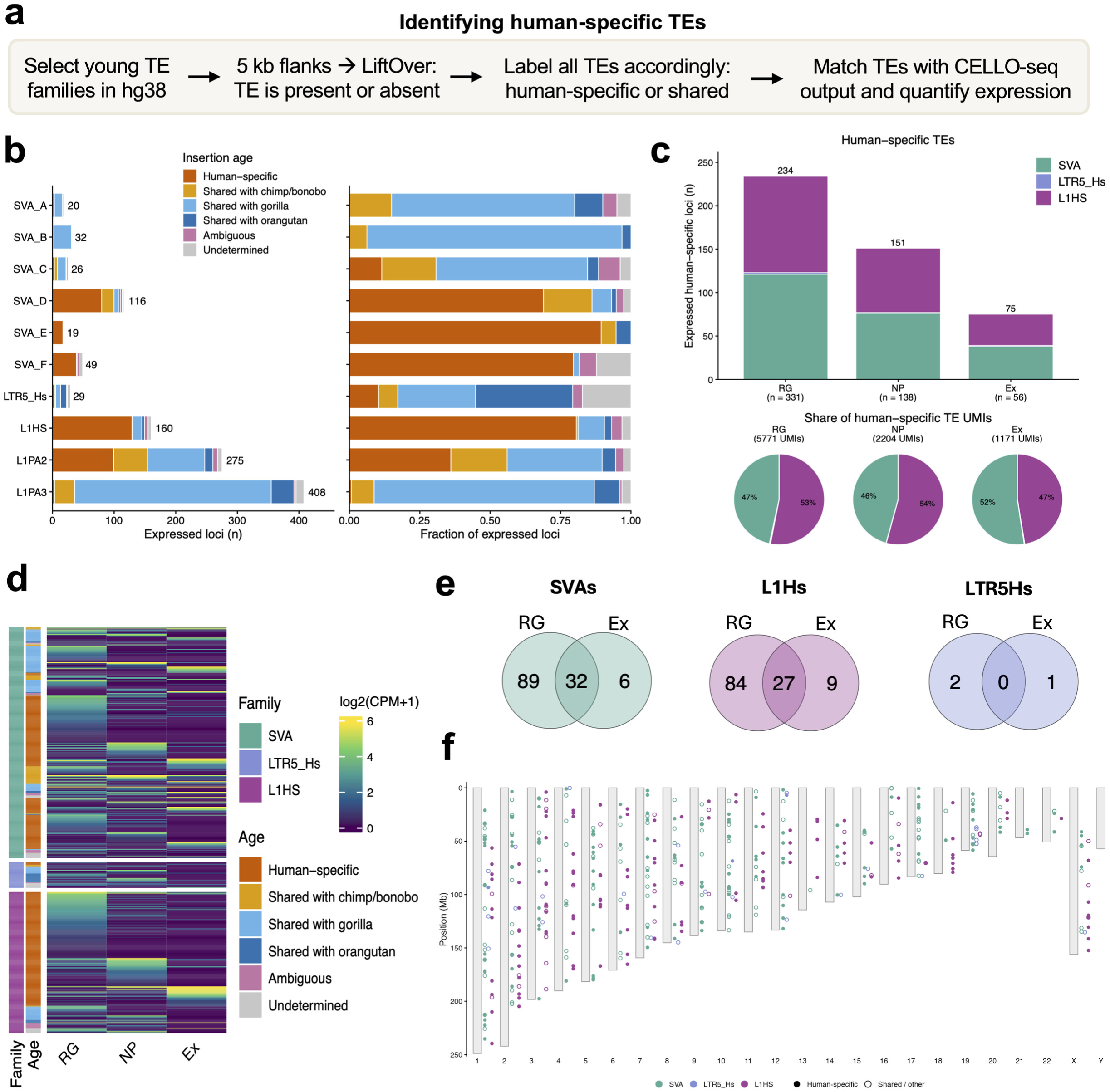
Cell-type-resolved distribution and evolutionary origin of expressed young transposable elements during early human corticogenesis. **(a)** Schematic of the pipeline used to identify human-specific TEs. **(b)** Left: bar chart showing the number of expressed loci per TE subfamily. Right: proportional contribution of each insertion age category to the expressed loci within each TE subfamily in d21 cortical organoids, highlighting the varying degrees of human-specificity across subfamilies, **(c)** Top: bar chart showing the total number of expressed human-specific TE loci per cell type. Bottom: pie charts showing the proportional contribution of SVA (teal) and L1Hs (pink) families to total human-specific TE UMIs in each cell type demonstrating a consistent family composition across cell types with L1Hs marginally dominant in RG and NP and SVAs marginally dominant in Ex. (d) Heatmap of log2(CPM+1) expression values for all expressed human-specific TE loci across RG, NP, and Ex cell types. The heatmap reveals both constitutively expressed loci shared across cell types and cell-type-restricted loci, with the majority of progenitor-restricted elements found in RG. (e) Venn diagrams showing the overlap of expressed human-specific TE loci between RG and Ex for SVAs, L1 Hs, and LTR5Hs, demonstrating the strong progenitor bias in TE locus expression, (f) Ideogram showing the chromosomal distribution of all expressed human-specific TE loci across the genome (human-specific: filled circles, shared/other: open circles). Expressed loci are distributed broadly across all autosomes and the X chromosome, with no obvious chromosomal clustering, consistent with the genome-wide regulatory potential of these elements.

To dissect the transcriptional activity of human-specific TEs at cellular resolution, we next mapped expressed loci to each cell type identified within the cortical organoids (**Fig. 3c,d**). Radial glia (n=331 cells) expressed the most human-specific TE loci (234 total), followed by neural progenitors (n=138 cells, 151 loci) and excitatory neurons (n=56 cells, 75 loci). The relative contribution of each subfamily to the expressed repertoire was remarkably consistent across cell types: SVA and L1Hs elements contributed approximately equally across all three populations, with L1Hs accounting for a marginally greater share in radial glia (53%) and neuronal progenitors (54%), while SVAs were marginally dominant in excitatory neurons (52%). LTR5Hs elements represented a minor fraction of total reads across all cell types (**Fig. 3c,d**), and no subfamily appeared to be preferentially silenced or activated during early neurogenesis. When examining locus-specific expression, 32% of SVAs (32/101) were constitutively expressed across radial glia and excitatory neurons, while the majority showed cell-type-specific expression, with progenitor-restricted loci outnumbering neuron-restricted ones approximately 14:1 (70% vs 5%, **Fig. 3e**). L1Hs showed a similar pattern, with 23% of loci shared between radial glia and excitatory neurons, 70% unique to radial glia, and 8% unique to excitatory neurons. The apparent bias toward progenitor-restricted expression should nonetheless be interpreted cautiously, as the unequal cell numbers across groups (RG n=331 vs Ex n=56) may inflate the number of loci detected in radial glia relative to more sparsely sampled populations. Overall, expressed human-specific TE loci are distributed across all autosomes and the X chromosome, suggesting genome-wide transcriptional activity (**Fig. 3f**).

Of note, we additionally detected the expression of young AluY subfamily members, the majority of which carried human-specific insertions (**Supp. Fig. 3**). However, Alu elements are transcribed by RNA polymerase III, which does not polyadenylate its products, whereas they carry a genomically encoded 3’ poly-A tract and an internal A-rich linker, both of which are efficiently bound by oligo-dT. Their recovery by poly-A-based CELLO-seq therefore cannot be taken as evidence of autonomous transcription, and may instead reflect internal priming or Alu sequence carried within polyadenylated Pol II transcripts. Therefore Alus were excluded from further analyses.

### SVAs are widely expressed across cell types during brain development and carry motifs for cortical and neurogenic transcription factors

Given that SVAs represent the youngest TE family exclusive to great apes, and have previously been implicated as cis-regulatory elements in hippocampal neurogenesis, neural crest development and pluripotent stem cells^12–14^, we next examined the chromatin accessibility of expressed SVAs across multiple timepoints of cortical organoid differentiation using bulk ATAC-seq data (**Fig. 4a**). SVA elements showed their greatest chromatin accessibility at d4 of differentiation, corresponding to the neuroectodermal transition, with overall accessibility declining over six months of differentiation (d181). This observation is consistent with the greater progenitor-restricted expression pattern observed by CELLO-seq and together suggesting their greatest regulatory influence may be concentrated at the earliest stages of cortical fate commitment (**Fig. 4a**).

**Figure 4.**
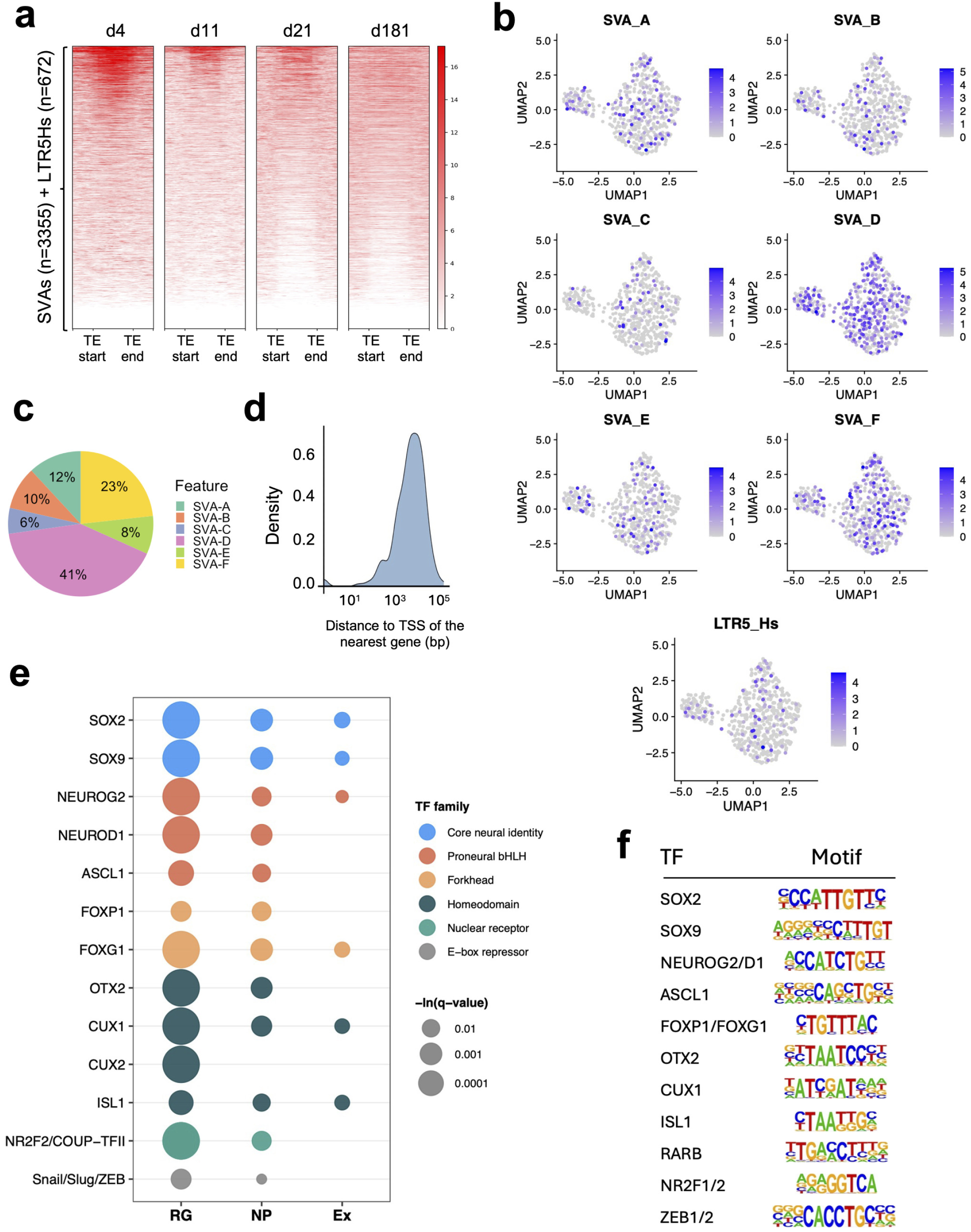
SVA chromatin accessibility dynamics across cortical organoid differentiation and transcription factor motif enrichment in expressed SVA-derived enhancers. **(a)** Heatmaps of bulk ATAC-seq signal across all SVA and LTR5Hs loci across four time points of cortical organoid differentiation show their greatest chromatin accessibility at d4. **(b)** UMAP feature plots showing pseudobulk ln(CP1 OK + 1) expression of each SVA subfamily and LTR5Hs in the CELLO-seq dataset, with SVA_D and SVA_F showing the highest expression levels, **(c)** Proportional expression levels of each SVA subfamily, **(d)** Density plot showing the distribution of distances (bp) from expressed SVA loci to the transcription start site (TSS) of their nearest gene. The distribution is strongly right-skewed, with the majority of expressed SVAs located more than 1 kb from their nearest gene TSS, consistent with a distal enhancer function, **(e)** Dot plot showing the enrichment of key TF binding motifs in expressed SVAs across radial glia (RG), neural progenitors (NP), and excitatory neurons (Ex), identified by HOMER motif enrichment analysis (all motifs q < 0.05). **(f)** Transcription factor binding motifs enriched in expressed SVAs compared to non-expressed SVAs across cell types.

At pseudobulk level, all SVA subtypes are uniformly expressed across cell types at d21, with SVA_D and SVA_F showing the highest expression (**Fig. 4b,c**). The majority of expressed SVAs are located more than 1kb from their nearest gene transcription start site, consistent with a distal enhancer function rather than direct promoter activity (**Fig. 4d**). To investigate whether these SVAs carry regulatory potential relevant to cortical development, we performed motif enrichment analysis using HOMER on expressed SVAs (CELLO-seq data) compared to non-expressed ones within each cell type. To confirm biological relevance, we verified that the transcription factors corresponding to enriched motifs are themselves expressed in the corresponding cell types (**Supp. Fig. 4**). The most consistently enriched motifs across all three cell types were those of the SOX family (including SOX2 and SOX9), reflecting the established role of SOX factors as master regulators of neural progenitor identity (**Fig. 4e,f)**. Proneural bHLH factors, including NEUROG2, NEUROD1, and ASCL1, were also enriched, alongside Forkhead family motifs (including FOXP1 and FOXG1) and homeodomain factor OTX2, consistent with their established role in cortical neuron identity. Homeodomain factors CUX1 and CUX2 were enriched in early excitatory neurons, consistent with early establishment of upper-layer cortical identity, while additional enriched motifs included ISL1 and RARβ, and E-box repressor motifs shared by the Snail/Slug/ZEB family were detected in radial glia and neural progenitors. An interesting finding was the enrichment of NR2F2 (COUP-TFII) motifs in radial glia and neural progenitors, given NR2F2’s established role in the regional as well as temporal patterning of cortical radial glia.

Together, these findings indicate that expressed SVAs carry a rich repertoire of binding sites for transcription factors governing neural progenitor identity, neuronal differentiation timing, and cortical patterning, positioning them as putative cis-regulatory elements embedded within the core transcriptional programmes of early human corticogenesis.

### Young TE silencing affects pathways related to neurogenesis

To test whether these TEs are functionally required for the transcriptional programmes their motifs suggest, we silenced both SVAs and LTR5Hs using an established CRISPRi model^20^ which targets a conserved region between both SVAs and LTR5Hs, derived from the same ancestral origin (HERVK-10), during cortical organoid differentiation (**Fig. 5a**). We found 589 accessible SVAs and LTR5Hs at d4 of differentiation, of which 90% showed significantly reduced accessibility following CRISPRi treatment (**Fig. 5b**). Single-cell transcriptomic analysis of d21 cortical organoids revealed widespread transcriptional dysregulation across all four major cell populations. In total, 654 differentially expressed genes were identified (DEGs, FDR <0.05), corresponding to 517 unique genes after accounting for those shared between cell types (**Fig. 5c**).

**Figure 5.**
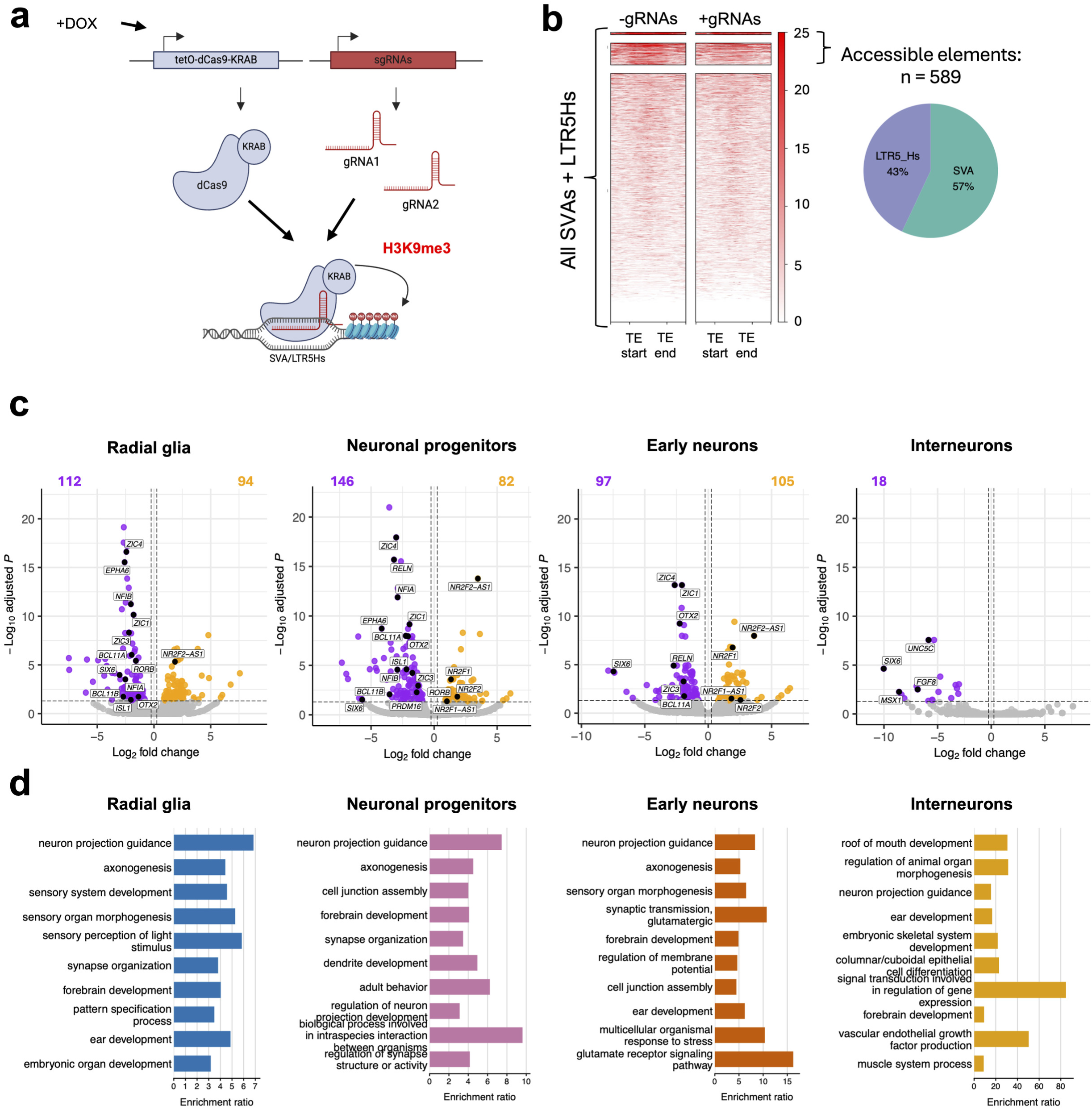
CRISPRi silencing of SVAs and LTR5Hs causes widespread transcriptional dysregulation across cortical cell types. **(a)** Schematic of the CRISPRi system used to silence SVAs and LTR5Hs. Adapted from Deelen et aL, 2025. **(b)** Left: heatmaps of ATAC-seq signal across all accessible SVA and LTR5H loci (n=589) in control (-gRNAs) and CRISPRi (÷gRNAs) organoids at d4, aligned from TE start to TE end. The majority of accessible elements show a marked reduction in chromatin accessibility upon CRISPRi treatment, with 90% of accessible elements less accessible in ÷gRNA cells. Right: pie chart showing the proportional composition of the 589 accessible elements, comprising SVAs (57%) and LTR5Hs (43%). (c) Volcano plots showing differential gene expression between CRISPRi (÷gRNAs) and control (-gRNAs) organoids at d21 for each of the four major cell types. Significantly downregulated genes (FDR < 0.05) are shown in purple, and significantly upregulated genes in orange, **(d)** Bar charts showing GO biological process enrichment analysis of downregulated DEGs for each cell type. Radial glia (blue), neural progenitors (pink), and early excitatory neurons (orange) converge on shared terms including neuron projection guidance, axonogenesis, forebrain development, and synapse organisation, consistent with broad impairment of cortical wiring and connectivity programmes. Excitatory neurons additionally show enrichment for synaptic transmission and glutamate receptor signalling terms. Interneurons (yellow) show a distinct GO profile dominated by terms related to morphogenesis and organ development.

One dominant theme was a loss of dorsal cortical identity, evidenced by consistent downregulation of *ZIC1/3/4*, *OTX2* and its antisense transcript *OTX2-AS1*, and *PRDM16* across radial glia and neuronal progenitors. Additionally, genes required for the correct layering and identity of cortical neurons were also downregulated. *RELN*, which encodes the extracellular matrix glycoprotein reelin and is essential for the inside-out patterning of cortical layers, was downregulated in both neural progenitors and excitatory neurons. The corticofugal fate specifiers *BCL11A* and *BCL11B* were suppressed in progenitors and radial glia respectively, as were the layer-identity determinants *RORB* (layer IV) and *ISL1*. The axon guidance receptor *EPHA6* was also significantly downregulated in both radial glia and neural progenitors. Beyond cortical identity and layering, *NFIA* and *NFIB*, two transcription factors that orchestrate gliogenesis and are required for the specification of upper-layer neurons and astrocyte precursors, were also concurrently suppressed. Early interneurons showed a qualitatively distinct response: all 18 DEGs were downregulated, with several showing large effect sizes, including *SIX6* (log2FC = −10.0), *MSX1* (−8.6), *FGF8* (−6.9), and *UNC5C* (−5.8), genes collectively implicated in interneuron patterning and fate specification, and tangential migration.

CRISPRi silencing of SVAs and LTR5Hs led to significant upregulation of *NR2F2-AS1* in radial glia, followed by additional upregulation of *NR2F2*, *NR2F1*, and *NR2F1-AS1* in neural progenitors and excitatory neurons, with a progressive increase in effect size across these loci suggesting an amplifying transcriptional cascade that propagates from radial glia through the neurogenic lineage (**Supp. Fig. 5**). This is consistent with the enrichment of *NR2F2* motifs in expressed SVAs, suggesting that SVA- or LTR5Hs-derived enhancers may normally act to repress this regional patterning axis.

Across radial glia, neural progenitors, and early excitatory neurons, gene ontology (GO) enrichment analysis of downregulated genes converged on shared terms including neuron projection guidance, axonogenesis, forebrain development, and synapse organisation, consistent with a broad impairment of cortical wiring and connectivity programmes (**Fig. 5d**). Enrichment of glutamate receptor signalling and synaptic transmission terms among excitatory neuron DEGs suggests functional consequences at the level of nascent synaptic identity. Taken together, these findings implicate SVA- and LTR5Hs-derived regulatory elements in maintaining a transcriptional programme of dorsal cortical identity, layer specification, and progenitor maturation during early human neurogenesis.

### The induced silencing of SVAs and LTR5Hs during brain organoid formation leads to a delay in radial glia-to-neuron differentiation

Single-cell RNA sequencing revealed a shift in cell type composition between CRISPRi (+gRNA) and control (-gRNA) organoids, with the +gRNA condition showing a marked enrichment of radial glia (∼80% vs ∼70%) and a corresponding reduction in neural progenitors (∼3.6% vs ∼4.5%), excitatory neurons (∼11% vs ∼17%) and interneurons (∼0.8% vs ∼2.8%), consistent with impaired progression through the neurogenic programme (**Fig. 6a)**. To determine whether this compositional shift reflects a delay in differentiation, we performed pseudotime trajectory analysis using Monocle3, which orders cells along a developmental path based on transcriptional similarity. This analysis confirmed a highly significant difference in pseudotime distribution between conditions: -gRNA cells had a median pseudotime of 25.6 compared to 15.7 in +gRNA cells (Wilcoxon test, balanced n: p < 2.2 × 10^-16^, **Fig. 6b**). Notably, at the earliest pseudotime values (0-15), the +gRNA condition retained approximately three times as many cells as the -gRNA condition, while at later pseudotime values (>20), two clear peaks, corresponding to more advanced differentiation stages were markedly reduced in the +gRNA condition, indicating that SVA and LTR5Hs silencing impairs progression through the intermediate and late stages of cortical differentiation.

**Figure 6.**
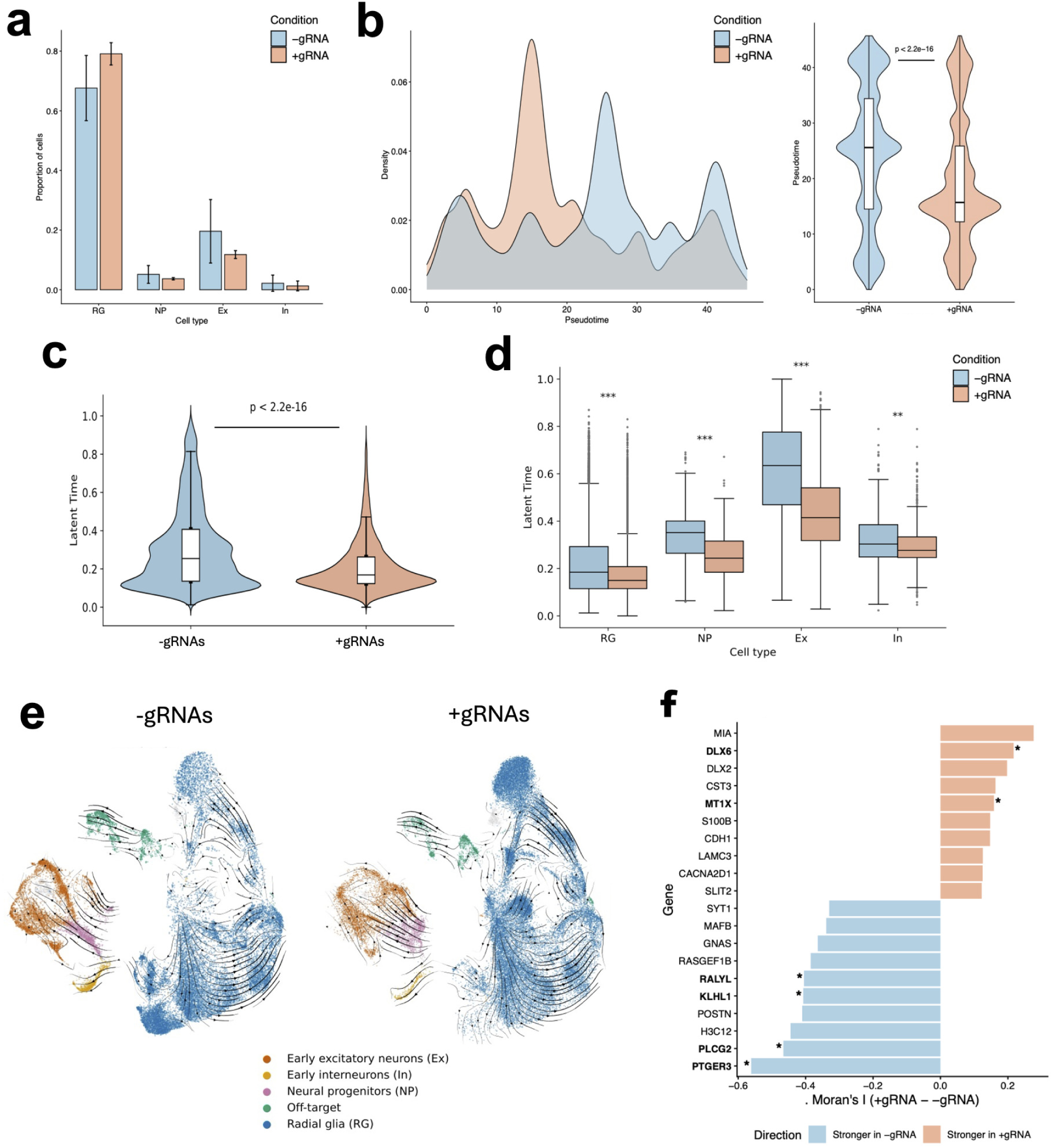
SVA and LTR5H silencing causes a delay and partial redirection of radial glia-to-neuron differentiation. **(a)** Bar chart showing the proportion of each cell type in CRISPRi (÷gRNA) and control (-gRNA) organoids at d21. CRISPRi organoids show a marked enrichment of RG (∼80% vs ∼70%) and a corresponding reduction in all neuronal populations. Error bars indicate standard deviation across replicates (n=2). (b) Density plots of Monocle3 pseudotime values for ÷gRNA and -gRNA cells. Control cells show a broader distribution with two prominent peaks at later pseudotime values (>20), corresponding to more advanced differentiation stages, while CRISPRi cells are enriched at earlier pseudotime values (0-15) and show a compressed distribution, indicating impaired progression through intermediate and late stages of cortical differentiation. Median pseudotime: -gRNA = 25.6, ÷gRNA = 15.7 (Wilcoxon test, p < 2.2 × 10"^16^). **(c)** Violin plots showing the distribution of scVelo latent time values across all cell types combined. CRISPRi cells show a significantly lower median latent time and a more compressed distribution enriched at early developmental stages (p < 2.2 × 10“^16^), independently confirming the differentiation delay observed by pseudotime analysis, **(d)** Box plots showing scVelo latent time values per cell type for ÷gRNA and -gRNA conditions. All four cell populations show significantly higher latent time in control cells (*** p < 0.001, ** p < 0.01), with the most pronounced difference observed in excitatory neurons, (e) RNA velocity stream plots for -gRNAs and ÷gRNAs organoids, with cells coloured by annotated cell type: early excitatory neurons (Ex, orange), early interneurons (In, yellow), neural progenitors (NP, pink), off-target (green), and radial glia (RG, blue). Velocity arrows flow from radial glia through neural progenitors to excitatory neurons in both conditions, indicating that the overall trajectory direction is conserved, (f) Bar chart showing genes with differential trajectory structure between conditions, quantified by Moran’s I spatial autocorrelation. Genes shown in blue have stronger trajectory structure in control (-gRNA) cells, including neuronal maturation markers PTGER3, PLCG2, KLHL1, RALYL, and SYT1, collectively implicated in synaptic vesicle release and cortical maturation. Genes shown in salmon have stronger trajectory structure in CRISPRi (÷gRNA) cells, including neural crest markers MIA and CDH1, ventral forebrain markers DLX2 and DLX6, and non-neuronal programme markers MT1X, SWOB, and CST3. Asterisks (*) denote genes that are also significantly differentially expressed between conditions.

Interestingly, we observed a circular RNA velocity loop within the radial glia subpopulation, which could indicate different stages or subtypes present within radial glia, which are known to show transcriptional heterogeneity^30^. Cell cycle scoring confirmed the overall direction of differentiation, with neuronal cells predominantly in G1 state (**Supp. Fig. 6**). The circular RNA velocity pattern observed within the RG cluster is consistent with active cell cycle progression: all three cell cycle phases are represented within the RG population in a spatially ordered fashion on the UMAP, reflecting the sequential transitions from S phase through G2M and back to G1 (**Supp. Fig. 6a**). Cell cycle scoring showed a significantly higher proportion of +gRNA cells in G1 phase compared to -gRNA cells (44.3% vs. 36.5%, χ² = 232.75, p < 2.2 × 10⁻^16^; **Supp. Fig. 6b**), suggesting impaired G1 exit and reduced commitment to either further proliferation or differentiation. This accumulation of RG cells at the G1 checkpoint is consistent with the broader pattern of progenitor retention observed across the pseudotime and latent time analyses, implicating SVAs and LTR5Hs in the transcriptional regulation of this critical cell fate transition.

To interrogate the transcriptional dynamics underlying the observed differentiation delay, we next performed RNA velocity analysis using scVelo, which infers a cell’s differentiation state (termed latent time) from the ratio of unspliced to spliced mRNA, which reflects the current direction and speed of transcriptional change. Latent time analysis confirmed that +gRNA cells have a significantly lower median latent time (p < 2.2 × 10⁻¹⁶) and a more compressed distribution with cells enriched at early developmental stages, in support of the proposed developmental delay (**Fig. 6c**). Critically, all cell populations showed significantly higher latent time values in -gRNA cells (all p < 0.001), with the most pronounced difference observed in excitatory neurons (**Fig. 6d**). This is consistent with the impaired progression through intermediate and late differentiation stages observed in the pseudotime analysis, and with the downregulation of glutamate receptor signalling and synaptic transmission pathways specifically in excitatory neurons in the differential gene expression analysis.

While the overall velocity trajectory direction flowing from radial glia through neural progenitors to excitatory neurons was conserved between conditions (**Fig. 6e**), the transcriptional content of the trajectory shifted significantly upon SVA and LTR5H silencing, as revealed by Moran’s I spatial autocorrelation analysis (**Fig. 6f**). In control cells, genes with the strongest trajectory structure were neuronal maturation markers including *PTGER3*, *SYT1*, *RALYL*, *KLHL1*, and *PLCG2*, collectively implicated in synaptic vesicle release and cortical maturation. In CRISPRi cells, this was replaced by genes associated with neural crest (*MIA*, *CDH1*), ventral forebrain (*DLX2*, *DLX6*), and non-neuronal programmes (*S100B*, *CST3*, *MT1X*). This shift suggests that SVA and LTR5H silencing not only delays but partially redirects the transcriptional character of differentiating cells toward non-cortical states, a shift that falls short of a complete fate change but represents a meaningful departure from canonical dorsal cortical identity.

### Co-expression network analysis reveals dysregulation of coordinated transcriptional programs across cortical cell types

We then performed the high-dimensional weighted gene co-expression network analysis (hdWGCNA) to investigate which co-expression modules are at play in brain organoids and which are disrupted upon SVA and LTR5Hs silencing. This identified 5-20 co-expression modules per cell type, each comprising a set of co-regulated genes with shared biological functions represented by their most highly connected members (hub genes) (**Fig. 7a,b**). Module activity was then compared between CRISPRi (+gRNAs) and control (-gRNAs) organoids to identify programmes systematically gained or lost upon TE silencing.

**Figure 7.**
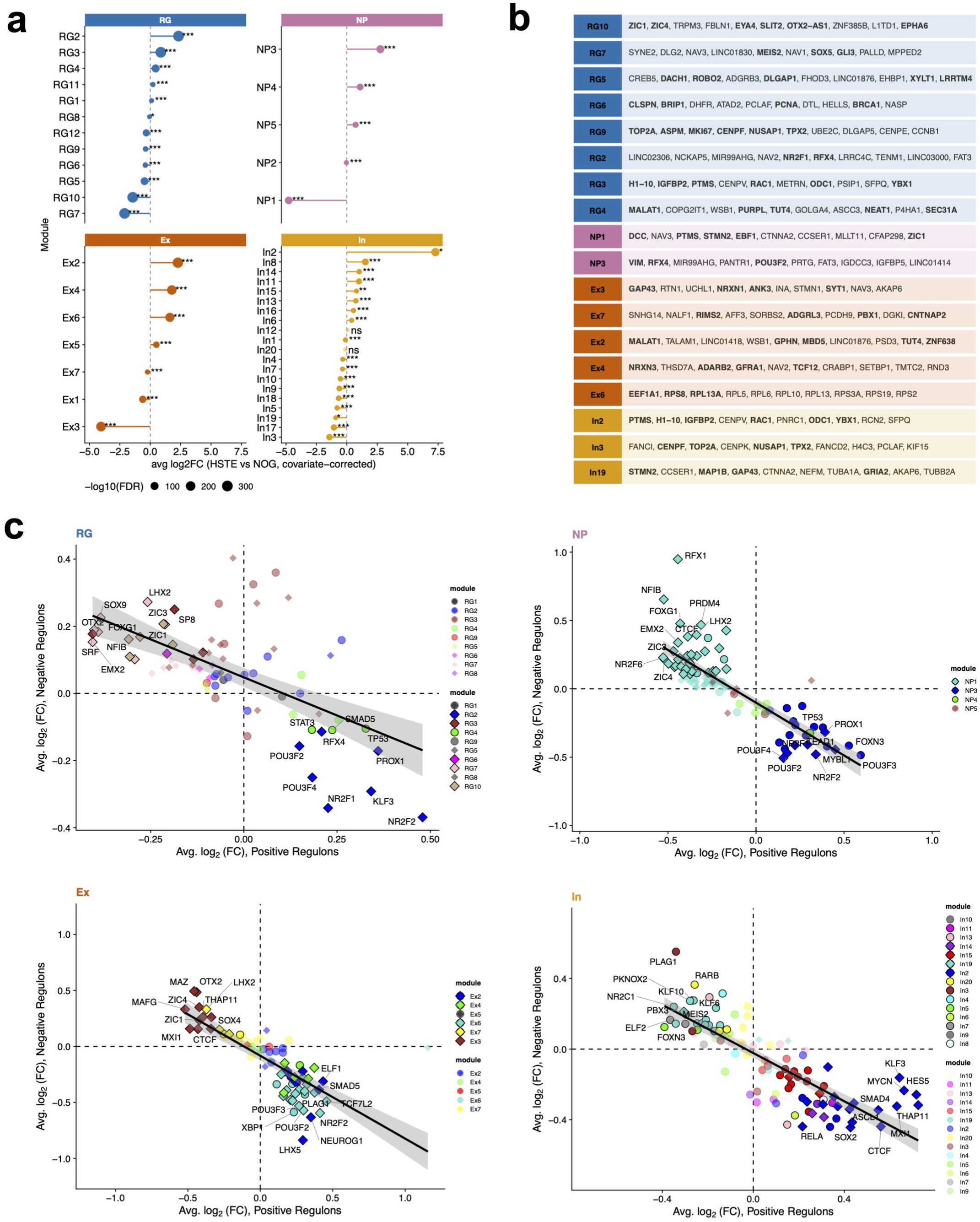
Co-expression network analysis reveals coordinated transcriptional programmes dysregulated by SVA and LTR5H silencing across cortical cell types. **(a)** Lollipop plots showing the average log2 fold change (÷gRNAs vs -gRNAs, covariate-corrected) of co-expression module activity for each cell type identified by hdWGCNA. Significance thresholds are indicated (*** FDR < 0.001, ns = not significant), **(b)** Table of the top ten hub genes for each highlighted module, with key genes shown in bold typeface, **(c)** Scatter plots showing regulon activity for radial glia (RG, top left), neural progenitors (NP, top right) excitatory neurons (Ex, bottom left) and interneurons (In, bottom right) in CRISPRi versus control organoids. Each point represents one transcription factor regulon, with the x-axis showing the average log2 fold change of positive regulon targets (genes activated by the TF) and the y-axis showing the average log2 fold change of negative regulon targets (genes repressed by the TF) in CRISPRi relative to control. Points are coloured by hdWGCNA module membership. The diagonal regression line indicates the overall regulon activity trend. TFs in the upper-left quadrant (positive targets down, negative targets up) represent lost activator regulons in CRISPRi. TFs in the lower-right quadrant (positive targets up, negative targets down) represent gained regulons in CRISPRi. Selected TFs are labelled.

In radial glia, the dominant pattern was loss of dorsal cortical identity. The two most significantly downregulated modules, RG10 and RG7, together define a dorsal cortical identity programme with RG10 hub genes including *ZIC1*, *ZIC4*, *OTX2-AS1*, *EPHA6, SLIT2* and *EYA4*, and RG7 hub genes including *MEIS2*, *GLI3*, and *SOX5*. Critically, 9 of 10 RG10 hub genes were independently identified as DEGs. Axon guidance (RG5), cell cycle (RG9) and DNA replication (RG6) modules were also downregulated, underscoring the breadth of identity loss. Against this background of lost dorsal identity, three modules were upregulated: a ventralisation module (RG2: *NR2F1*, *RFX4*), a progenitor maintenance module (RG3), and a stress/lncRNA module (RG4), collectively describing radial glia shifted toward an earlier, less differentiated, and more ventrally specified state. A similar pattern was evident in neural progenitors, where the neuronal differentiation module NP1 (*ZIC1*, *EBF1*, *DCC, STMN2*; 6/10 hubs as DEGs) was the most significantly downregulated, while the progenitor identity module NP3 (*VIM*, *RFX4*, *POU3F2*) was upregulated, indicating impaired exit from the progenitor state. For excitatory neurons, pan-neuronal identity module Ex3 (*GAP43*, *NRXN1*, *SYT1*, *ANK3*) was the most significantly downregulated, alongside mature synaptic module Ex7, which contains ASD-risk genes including *CNTNAP2*, *ADGRL3*, and *RIMS2*. Three modules were upregulated: ribosomal activity (Ex6), stress/lncRNA (Ex2), and early synaptic adhesion (Ex4: *NRXN3*, *ADARB2*, *GFRA1*). The simultaneous downregulation of mature synaptic programmes and upregulation of earlier-acting synaptic identity genes suggests that CRISPRi excitatory neurons are arrested at an immature synaptic state, unable to progress to mature connectivity. In interneurons, the module data revealed a phenotype not apparent from the DEG analysis alone. In2, a progenitor maintenance module (*PTMS, H1-10, IGFBP2, RAC1, YBX1*), had the largest effect size in the entire dataset (log2FC = 7.3). In contrast, mature neuronal identity (In19: *STMN2*, *GAP43, MAP1B, GRIA2*) and cell cycle (In3) modules were all significantly downregulated. This combination suggests that CRISPRi interneurons are arrested in a non-proliferating, fate-indeterminate progenitor state. This differentiation block parallels the G1 arrest in radial glia and points to a shared mechanism of impaired neurogenic exit, despite their distinct developmental origins.

Complementing the module activity analysis, regulon inference identified consistent upstream drivers across cell types (**Fig. 7c**). Across radial glia, neural progenitors, and excitatory neurons, the consistently lost regulons (SOX9, OTX2, FOXG1, EMX2, LHX2, NFIB, and ZIC1/3/4) map directly onto the downregulated dorsal identity modules RG10, RG7, and NP1, identifying these TFs as the likely mediators of the identity loss programme. Moreover, the consistently gained regulons (NR2F1/2 and POU3F2/3) map onto the upregulated ventralisation (RG2, NP3) modules respectively. In interneurons, the most strongly upregulated regulons, HES5, ASCL1, MYCN, and SMAD4 all map to the In2 progenitor maintenance module and collectively implicate aberrant progenitor maintenance signalling as a driver of the differentiation block, while downregulated regulons included MEIS2 and RARB, consistent with loss of dorsal forebrain patterning and retinoic acid signalling in this population. Strikingly, several TFs whose binding motifs are enriched in expressed SVAs and LTR5Hs are themselves among the most significantly dysregulated regulons upon CRISPRi. Among those lost, SOX9, OTX2, FOXG1, FOXP1, CUX1, MEIS2, GLI3, ZEB1, and RARβ all appear as downregulated regulons. These map directly onto the downregulated dorsal identity modules RG7, RG10, and NP1, reinforcing their role as mediators of the identity loss programme. Notably, MEIS2 and GLI3 are additionally serving as hub genes of RG7. Together, these results provide three-layer convergence between motif content, regulon activity, and co-expression module structure. Among those gained, NR2F1/2 motifs enriched in RG and NP SVAs/LTR5Hs correspond to the most consistently upregulated regulons across all three cell types, while ASCL1 motifs correspond to an upregulated regulon in the interneuron progenitor maintenance module In2.

Together, these convergences directly link the chromatin-level motif landscape of expressed SVAs to the transcriptional consequences of their silencing, supporting a model in which SVA and LTR5H-derived enhancers serve as binding platforms for dorsal cortical identity TFs. Their loss propagates dysregulation across the neurogenic programme, manifesting as progenitor retention, impaired G1 exit, transcriptional ventralisation, and fate-indeterminate arrest in interneuron precursors. Collectively, these findings implicate SVAs and LTR5Hs as functional components of the human-specific regulatory architecture governing the timing of early cortical neurogenesis.

## Discussion

In this study, we applied locus-resolved single-cell long-read sequencing to characterise the expression and regulatory potential of young human-specific TEs during early cortical neurogenesis, and used CRISPRi to directly test their functional requirement. We establish the expression of tens of individual loci from human-specific members of multiple TE subfamilies including SVAs, LTR5Hs and L1Hs. While the expression of SVAs during brain development has been investigated previously^12,20^, their endogenous locus-specific expression of individual members has only been investigated in specific functional contexts.

For LTR5Hs, the literature is even more limited. While HERVK (HML-2) activity has been shown to influence cortical neuron differentiation and forebrain patterning^32^, LTR5Hs, the most evolutionarily recent and human-enriched HERVK subfamily, has not been characterised in terms of its expression or regulatory activity during brain development at any level of resolution to our knowledge. Evolutionarily young LINE-1 subfamilies have received more attention, with recent studies demonstrating their expression and functional roles during human cortical neurogenesis, and individual L1Hs insertions have been implicated in specific genomic and disease contexts^33–35^. Yet, none have resolved their expression systematically at individual locus resolution using long-read sequencing. Our study therefore provides the first locus-resolved transcriptional atlas of all three families simultaneously in a human cortical single-cell context.

We demonstrate that human-specific SVA and LTR5Hs expressed during early corticogenesis carry binding motifs for dorsal cortical identity transcription factors, and that their silencing causes a coordinated failure of the radial glia-to-neuron transition characterised by progenitor retention, transcriptional ventralisation, and impaired synaptic maturation. The convergence of motif enrichment, regulon inference, and co-expression network analysis points to a model in which SVA-derived enhancers normally serve as binding platforms for TFs including FOXG1, OTX2, SOX9, NFIB, and NR2F2, whose combined activity sustains the dorsal cortical identity and neurogenic exit programmes of radial glia. Upon silencing, the loss of these binding platforms propagates transcriptional disruption across co-expression modules governing dorsal identity, cell cycle exit, and neuronal differentiation. We note that this model is based on motif enrichment and regulon activity and that ChIP-seq or CUT&RUN for these TFs in control versus CRISPRi cells would validate these findings mechanistically.

Notably, OTX2 motifs were enriched in radial glia and excitatory neuron SVAs, and *OTX2* was among the most significantly downregulated genes upon CRISPRi. Together, this is consistent with a model in which SVA-derived enhancers carrying OTX2 binding sites normally sustain *OTX2* expression and its regulatory function, and their silencing leads to loss of anterior cortical identity. The nearest SVAs and LTR5Hs lie over 1 Mb upstream or 350 kb (SVA_D) downstream of the *OTX2* gene, suggesting either long-range enhancer activity or the *OTX2* downregulation is an indirect downstream consequence of silencing other SVA-regulated targets. Future chromatin conformation capture experiments and targeted perturbation of the downstream SVA_D elements in radial glia could directly test whether these elements physically contact the OTX2 promoter and are required for its expression.

Another remarkable finding is the upregulation of *NR2F2*, *NR2F2*-*AS1*, *NR2F1*, and *NR2F1*-*AS1* upon CRISPRi, with a progressive increase in effect size across these loci suggesting an amplifying transcriptional cascade. NR2F2 is a key temporal patterning factor in cortical radial glia, normally acting to suppress early cortical identity and promote the sequential production of later-born neuronal subtypes^36^. Its upregulation upon SVA/LTR5Hs silencing is consistent with a premature temporal identity shift that would compound the ventralisation phenotype. Notably, *NR2F2-AS1* itself harbours an intronic LTR5Hs element, raising the possibility that this element normally represses *NR2F2-AS1* transcription, and that its silencing by CRISPRi derepresses *NR2F2-AS1*, which in turn cis-activates *NR2F2* sense transcription. This model would account for the progressive amplification of effect size observed across this locus and implicates an intronic LTR5Hs as a direct cis-regulator of NR2F2 temporal identity programmes. This provides a mechanistic precedent for how intronic TE insertions can alter the expression of dosage-sensitive neurodevelopmental regulators, with potential relevance to conditions associated with NR2F2 haploinsufficiency including intellectual disability and autism, and positions human-specific TEs as a previously underappreciated source of regulatory variation at the intersection of cortical development and neurological disease susceptibility, motivating locus-resolved investigation of human-specific TE variation in patient-derived organoid models.

The differentiation phenotype we observe is best described as a combination of delay and partial redirection. While the overall velocity trajectory direction is conserved, Moran’s I analysis reveals that young TE silencing shifts the transcriptional character of differentiating cells toward neural crest and ventral forebrain programmes. This finding carries interesting evolutionary implications: if SVAs and LTR5Hs normally sharpen and accelerate the radial glia-to-neuron transition in a human-specific manner, their emergence in the human lineage may have contributed to the precise timing of neurogenic exit that underlies cortical expansion. Extended radial glia identity has been proposed as one mechanism of cortical expansion in humans^37,38^, and our data suggest that SVA regulatory activity may normally constrain this window, a hypothesis directly testable by comparative CELLO-seq analysis in chimpanzee or gorilla organoids, where these elements are absent or less abundant. The chromatin accessibility data showing that SVA and LTR5Hs elements reach peak accessibility at d4 of differentiation, corresponding to the neuroectodermal transition, further motivates CELLO-seq at this earlier time point, particularly given reports of human-specific regulatory divergence at the neuroectoderm-to-radial glia transition^25,39^. Characterising the locus-resolved TE expression landscape at d4 would establish whether the regulatory potential we infer from motif analysis is already deployed at the moment of cortical fate commitment.

The interneuron differentiation block, reflected in the massive upregulation of the progenitor maintenance module In2 alongside loss of mature interneuron identity modules, is an interesting finding given that cortical interneurons conventionally originate from the ventral ganglionic eminence^40^. However, studies in primates have demonstrated that a subset of cortical interneurons can arise from dorsal progenitors^41–43^. Therefore, in these organoids, interneurons may derive from a transiently more ventralised population or from the dorsal progenitor pool. In case of the latter, these interneurons would be directly susceptible to disruption of dorsal progenitor programmes. However, secondary effects from failing radial glia altering the signalling environment for migrating interneuron precursors, cannot be excluded^44^. Resolving this will require cell-type-specific perturbation approaches beyond the scope of the current study.

Beyond their developmental roles, the findings presented here have direct implications for understanding the contribution of TE variation to human neurodevelopmental and neurological disease. Whole-genome sequencing and mobile element scanning studies have identified thousands of polymorphic TE loci, a subset of which act as cis-eQTLs affecting gene expression in the brain^45–50^. TE variation has been linked to disease susceptibility through alterations in chromatin state, splicing, transcript structure, and gene dosage^49,51–54^. Our data suggest that SVA and LTR5Hs elements are functionally required for the expression of a set of well-established neurodevelopmental disease genes during cortical neurogenesis. Among the downregulated DEGs upon CRISPRi, we identify *BCL11A^55^* and *BCL11B^56^* (intellectual disability and ADHD), *NRXN1^57^* and *CNTNAP2^58^* (autism spectrum disorder) and *RELN^59^*(lissencephaly and cortical malformation), with the strongest effects observed in neural progenitors and radial glia. The consistent downregulation of these genes across multiple cell types, alongside the broader disruption of neuron projection guidance, forebrain development, and synaptic organisation programmes, suggests that SVA and LTR5Hs regulatory activity during early corticogenesis is required to maintain expression of gene networks whose disruption underlies a range of neurodevelopmental conditions. Naturally occurring variation at individual TE loci could therefore subtly perturb these same programmes, contributing to phenotypic diversity in cortical development and risk for neurodevelopmental disorders, a hypothesis directly testable by integrating the locus-resolved TE atlas generated here with population-level genomic variation data.

## Limitations of the study

Whilst revealing locus-specific TE expression levels in the developing brain, the CRISPRi approaches silences all SVAs and LTR5Hs elements simultaneously, which leaves individual locus contributions unresolved. This limitation motivates locus-specific follow-up experiments using individual SVA deletions or targeted activation of specific loci of interest. Furthermore, we included TE expression of elements located in intronic regions, which may capture some readthrough transcription from upstream Pol II loci rather than autonomous TE transcription. However, we propose this to be a minority. As we performed CELLO-seq on our CRISPRi and control line (which share the same genetic background), there may be de novo TE insertions present in our specific line that are not represented in the hg38 reference genome. While these private insertions are unlikely to substantially alter our conclusions, their potential regulatory contributions at individual loci cannot be fully excluded. Finally, cortical organoids do not fully recapitulate in vivo neurogenesis, particularly for interneurons, and validation in primary human tissue or comparative great ape models will be important to establish the physiological and evolutionary relevance of these findings.

## Materials and Methods

### Cortical organoid generation

Cortical organoids were generated from human induced pluripotent stem cells (hiPSCs) engineered with an inducible CRISPRi system (see CRISPRi cell line) following an adapted version of the guided cortical spheroid protocol^27^. Briefly, hiPSCs were maintained on Geltrex-coated plates in mTeSR1 medium and passaged using EDTA. For organoid induction, embryoid bodies were formed from single-cell hiPSC suspensions at a density of 10,000 cells/well in slit-well 96-well plates in E6 medium supplemented with SB431542 (10 µM), dorsomorphin (2.5 µM), and XAV939 (2.5 µM), for days 1-5 to promote dorsal forebrain identity, with daily media changes of 65% media. From days 6-24, organoids were transferred to Neurobasal-A medium supplemented with B27 without vitamin A, EGF (20 ng/ml), and FGF2 (20 ng/ml) to expand neural progenitor populations (65% media changes). From days 16-24 the media was changed every other day. From days 25–42, medium was changed to Neurobasal-A with B27 without vitamin A, BDNF (20 ng/ml), and NT3 (20 ng/ml) to promote neuronal differentiation (65% media changes). From day 43 onwards, organoids were maintained in Neurobasal-A with B27 without vitamin A for maturation (75% medium changes). Doxycycline (2 µg/ml) was added from day 1 and maintained throughout differentiation to induce dCas9-KRAB expression. All organoids were cultured in slit-well plates allowing shared media between wells to increase reproducibility, as described in Bertucci et al. (2026)^27^. Organoid size was monitored at regular intervals by brightfield microscopy and measured using Fiji. Quality control was performed at 3 weeks, 2 months and 4 months of differentiation by RT-qPCR and immunohistochemistry for timepoint-specific markers.

### CRISPRi cell line

The inducible CRISPRi hiPSC lines used in this study were generated as previously described^12,14,20^. Briefly, the C2277 hiPSC line was co-transfected with a piggyBac-based plasmid encoding dCas9-KRAB under a tetracycline-inducible (tetO) promoter (p-dCas9-KRAB) and a piggyBac transposase plasmid (p-PB-Transposase, Systems Bioscience), conferring constitutive puromycin resistance for stable clone selection. For the +gRNAs line (C2277-B), a third piggyBac plasmid (p-sgRNA, Systems Bioscience) encoding two sgRNAs targeting a conserved sequence shared between SVA and LTR5Hs elements, targeting ∼80% of annotated copies in the human genome, and conferring geneticin resistance was additionally co-transfected. The -gRNAs line (C2277-A) received p-dCas9-KRAB and p-PB-Transposase only and serves as a negative control. Stable clones were selected using puromycin (0.125 μg/mL) alone (-gRNAs) or combined with geneticin (100 μg/mL) (+gRNAs), and both lines were authenticated by STR profiling and screened for mycoplasma contamination prior to use.

### 10X Genomics scRNA-seq library preparation

For scRNA-seq, two independent batches of five pooled d21 organoids per line (-/+gRNAs) were dissociated into single-cell suspensions by incubation in Papain for 120 minutes at 37°C with gentle agitation (300 rpm) and periodic trituration using a glass Pasteur pipette. Cells were pelleted by centrifugation, resuspended in ice-cold PBS, and passed through a 40 µm cell strainer. Cell viability and concentration were assessed by Trypan Blue exclusion using the Countess automated cell counter and were >90% viable. Samples were submitted to the Imperial Genomics Facility for scRNAseq. Single-cell libraries were prepared using the 10x Genomics Chromium platform with the Chromium Single Cell 3’ v4 kit according to the manufacturer’s instructions. Libraries were sequenced on an Illumina NovaSeq X.

### 10X Genomics scRNA-seq data processing and analysis

For scRNA-seq, raw sequencing data were processed using Cell Ranger v8.0.1 (batch 1) and v9.0.1 (batch 2) with alignment to the GRCh38 reference genome. Downstream analysis was performed in R using Seurat (v5.6.0). Cells were filtered to retain those with 200-6,000 detected genes, >1,000 UMI counts, and <5% mitochondrial reads. Count data were log-normalised and scaled with regression of sequencing depth and mitochondrial percentage. The top 3,000 highly variable genes, identified using variance-stabilising transformation (VST), were used for dimensionality reduction (PCA) and clustering (Louvain algorithm). Batch correction was performed using Harmony integration (v2.0.5). Cell type annotation was performed by comparing cluster marker genes against published single-cell reference datasets from the human neocortex at gestational week 17-18 and gestational week 6-10^28,30^. Off-target populations including neural crest cells (SOX10+) and placode cells (SIX1+) were identified and excluded from downstream analyses.

### Pseudotime trajectory analysis

Pseudotime analysis was performed using Monocle3^60^ (v1.4.26) (https://cole-trapnell-lab.github.io/monocle3/). Cells were ordered along a principal graph rooted in the most outer right nodes within the radial glia cluster based on expression of radial glia marker genes. Pseudotime values were compared between +gRNA and -gRNA conditions using a Wilcoxon rank-sum test with subsampling to equal cell numbers per condition to account for the compositional imbalance between conditions.

### RNA velocity analysis

RNA velocity analysis was performed using scVelo (v0.3.4). Spliced and unspliced mRNA counts were quantified using STARSolo’s Velocyto. RNA velocity was estimated using the dynamical model from which velocity confidence, latent time, and velocity pseudotime were derived as well as the velocity streams per sample. Latent time distributions were compared between conditions using a two-sided Mann-Whitney U test, applied both globally and per cell type, after downsampling to equal cell numbers. Moran’s I spatial autocorrelation analysis was performed using scVelo’s built-in function to identify genes with differential trajectory structure between conditions.

### Pseudobulk differential expression analysis

Differential expression analysis between CRISPRi (+gRNA) and control (-gRNA) conditions was performed using a pseudobulk approach. For each cell type, raw counts were summed across all cells from the same sample to generate pseudobulk count matrices, with genes absent across all cells removed. Differential expression was tested using edgeR’s (v.4.6.3) likelihood ratio test (LRT) with a negative binomial generalised linear model and TMM normalisation, using the design formula ∼ batch + condition, with +gRNAs as the coefficient of interest. Analysis was performed on four cell types: radial glia (RG), neural progenitors (NP), excitatory neurons (Ex), and interneurons (In). Genes with an adjusted p-value < 0.05 (Benjamini-Hochberg correction) were considered differentially expressed. GO enrichment analysis of DEGs was performed using WebGestalt^61^ against the GO Biological Process database.

### Cell cycle scoring

Cell cycle phase was assigned to individual cells using Seurat’s CellCycleScoring function. The proportion of cells in each cell cycle phase was compared between +gRNA and -gRNA conditions within the radial glia population using a chi-squared test.

### hdWGCNA co-expression network analysis

High-dimensional weighted gene co-expression network analysis was performed using hdWGCNA (v0.4.12) as described in Morabito et al. (2023)^62^. Genes expressed in at least 5% of cells were selected for analysis. Metacells were constructed using k=20 nearest neighbours within each cell type and sample, with a maximum of 5 shared cells between metacells, using the Harmony-corrected embedding. Co-expression networks were constructed independently for each cell type (RG, NP, Ex, In) using a signed network topology, with soft-power thresholds selected per cell type based on scale-free topology criteria. Module eigengenes were computed with sample as a grouping covariate. Module connectivity (kME) was calculated per cell type, and hub genes were defined as the top 10 genes per module ranked by kME. Module expression scores were computed using both the Seurat and UCell methods across the top 25 hub genes per module. To compare module activity between CRISPRi (+gRNA) and control (-gRNA) conditions, a covariate-corrected differential module expression (DME) approach. Technical covariates (library size, number of detected features, mitochondrial read percentage, and ribosomal read percentage) were regressed out from module eigengenes using a linear model, and a Wilcoxon test was applied to the residuals to identify differentially expressed modules (DMEs) between conditions. P-values were corrected using the Benjamini-Hochberg method (FDR < 0.05). Transcription factor regulatory network analysis was performed using hdWGCNA’s built-in TF network module. TF motifs were scanned against gene promoters using JASPAR2020 (CORE vertebrate collection) and motifmatchr, with genomic coordinates from hg38/EnsDb.Hsapiens.v86. TF regulatory networks were constructed using XGBoost-based regression, and regulons were assigned using top 10 TFs per gene with a regulatory threshold of 0.01. Differential regulon activity between +gRNAs and -gRNAs conditions was assessed per cell type using FindDifferentialRegulons.

### CELLO-sequencing library preparation

CELLO-seq was performed on five pooled d21 cortical organoids from the -gRNA control line. The pooled organoids were dissociated into single-cell suspensions by incubation in Papain for 120 minutes at 37°C with gentle agitation (300 rpm) and periodic trituration using a glass Pasteur pipette. Cells were pelleted by centrifugation, resuspended in ice-cold PBS, and passed through a 40 µm cell strainer. Cell viability and concentration were assessed by Trypan Blue exclusion using the Countess automated cell counter and were >90% viable. Single live cells were sorted into individual wells of three 384-well plate (BioRad, HSP3801) containing 1 µl of lysis buffer, using a MA900 Cell Sorter (Sony) at the WIMM Flow Cytometry Facility and snap frozen on dry ice. CELLO-seq was performed as described in Marlow et al. 2025 ^23^, with minor modifications and reagent volumes per well scaled to 0.5x to account for the reduced well size of a 384-well plate compared to a 96-well plate as previously used. Briefly, dT-oligonucleotides (IDT) were annealed to polyadenylated RNA in the presence of RNase inhibitor (ThermoFisher, EO0381) and reverse transcription was performed using template switch oligos (IDT), Maxima H Minus Reverse Transcriptase (ThermoFisher, EP0753) and ET SSB (NEB, M2401S). Excess dT oligos were digested with Exonuclease I (NEB, M0293L) before splint oligonucleotides (IDT) were ligated onto cDNAs using T4 ligase (M0202L). Excess proteins were digested with proteinase K (Takara, 639270). An AMpure XP (Beckman Coulter) bead cleanup was then performed at 0.5x ratio before PCR amplification with Terra Direct PCR polymerase (Takara, 639270) and Nextera R1 and R2 primers (IDT) in a total reaction of volume of 14 µl with 26 cycles. A post-PCR 0.5x bead cleanup in the 384 well plate was then performed, and the four quadrants of the plate were separately pooled into individual DNA LoBind 1.5 ml tubes (Fisher, 10051232) before a 0.5x clean-up was carried within the 1.5 ml tubes and eluting in a final volume of 100 µl nuclease-free water (ThermoFisher, 10977-049).

To assess cDNA concentration and length, High Sensitivity Qubit quantification (ThermoFisher, Q33231) and High Sensitivity Bioanalyser (Agilent, 5067-4626) was carried out on the four pooled quadrants of the 384-well plate. 3 additional PCR cycles were conducted with Terra Direct PCR polymerase (Takara, 639270) and Nextera R1 and R2 primers (IDT) by splitting the pooled 100 µl quadrant across eight PCR tubes (VWR, BR781332) and performing each PCR amplification in a 30 µl total reaction volume. Following PCR, the quadrants were re-pooled and subjected to an additional 0.5x AMPure XP bead cleanup and another round cDNA quality control via High Sensitivity Qubit and High Sensitivity Bioanalyser. Taking 200 fmol of DNA, each quadrant was then barcoded with the Oxford Nanopore Technologies Native Barcoding Kit 96 V14 (SQK-NBD114.96) with a separate native barcode according to manufacturer’s instructions, allowing for multiplexing of four 384-well quadrants onto a single PromethION flow cell for sequencing with a PromethION 2 Solo.

### CELLO-sequencing data processing and analysis

Outputted reads were live basecalled using the super high accuracy model (software version 7.9.8+dcae65e9f) and native barcodes were demultiplexed (Dorado version 7.9.8). The resulting fastq files were subject to the Sarlacc CELLO-seq analysis pipeline, described in Marlow et al. 2025. Briefly, internal adaptor quality control was conducted before demultiplexing to generate individual fastq files for each CELLO-seq cell barcode. Reads were filtered out reads containing internal adaptors and then were aligned to the GENCODE human GRCh38_p13 assembly with Minimap2. Reads were grouped by unique molecular identifier (UMI) and error correction of reads based on UMI. Flair (v2.0.0) was then used to assign TE as well as isoform counts (hg38), using the flair align, correct, collapse and quantify modules to generate a final counts table before downstream analysis in R. All FLAIR steps were run on concatenated FASTQ files pooled across all samples prior to collapse, with per-sample quantification performed afterwards. From the flair collapse output, each read was classified as genic (containing an Ensembl gene ID), repeat (overlapping a TE locus at the 5’ end of the transcript), repeat isoform (a novel TE-initiated isoform), internal TE (a TE overlapping a genic read), novel isoform (a genic read without a known transcript ID), or “other”. For reads spanning multiple TE loci, the 5’ end-most TE was retained given the strand orientation of CELLO-seq reads (first TE for minus-strand reads, last TE for plus-strand reads). Counts from the flair quantify output were then aggregated per locus identifier. Low-quality cells were removed using quickPerCellQC from scater, based on library size and number of detected features. Non-expressed features were removed, and counts were normalised using logNormCounts from scran (library-size normalisation followed by log2-transformation). After quality control visualisation, data were log-normalised, scaled, and dimensionality reduction was performed by PCA followed by Harmony integration across batches. Cell type labels were assigned using the same cluster annotation strategy supported by reference atlases as the 10X scRNAseq dataset.

### Identification of human-specific transposable elements

To identify human-specific TE insertions, we analysed five TE subfamilies known to contain recently active or human-specific members: SVA_A through SVA_F, LTR5Hs, L1HS, L1PA2, and L1PA3. For each element in hg38, RepeatMasker fragments within 1,000 bp were merged into single loci, and elements shorter than family-specific minimum lengths (L1: 500 bp, SVA: 500 bp, LTR5Hs: 300 bp, Alu: 150 bp) were excluded as likely remnants. 5 kb flanking sequences on either side of each insertion were lifted over to the chimpanzee (panTro6), gorilla (gorGor6), and orangutan (ponAbe3) genomes using the UCSC liftOver tool with a minimum mapping ratio of 0.3. If both flanks mapped to the same chromosome and strand with a gap ≤ 25% of the element length, the element was classified as absent in that species; if the gap was ≥ 60% of the element length, it was classified as present, allowing for partial insertions or mapping imprecision at flank boundaries. Elements were classified as human-specific, shared with chimpanzee/bonobo, shared with gorilla, shared with orangutan, ambiguous, or undetermined based on their presence/absence across all four outgroup genomes. Expressed TE loci were defined as loci with ≥1 total UMI across all cells and detected in ≥1 cell. TE locus-level counts from CELLO-seq were matched to the age annotation by genomic overlap, requiring at least 50% overlap of the shorter interval. Pseudobulk counts per cell type were computed by summing raw UMIs across cells of each type and normalised to counts per million (CPM) using the total RNA library size (nCount_RNA) as the denominator. For visualisation, CPM values were log2-transformed (log2(CPM+1)).

### Motif enrichment analysis

Motif enrichment analysis within each cell type was performed using HOMER^63^ on expressed SVA loci identified in the CELLO-seq data compared to non-expressed SVA loci with a comparable subfamily composition as background, using the findMotifsGenome.pl function with default parameters. Motifs with a Benjamini-Hochberg adjusted p-value < 0.05 were considered significantly enriched.

### Bulk ATAC-seq library preparation and data analysis

Bulk ATAC-seq was performed on cortical organoids at days 4, 11, 21, and 181 of differentiation using the ATAC-seq kit (Active Motif, 53150) according to manufacturer’s instructions. Briefly, 100,000 cells per sample were lysed in cold lysis buffer and nuclei were tagmented using Tn5 transposase for 30 mins at 37°C at 800 rpm. Tagmented DNA was purified and PCR-amplified. Libraries were assessed for size distribution using the TapeStation 2200 and sequenced on an Illumina NovaSeq X Plus to a depth of 60M reads per sample. Raw reads were trimmed using TrimGalore! and aligned to hg38 using using the Burrows-Wheeler Alignment tool, with the MEM algorithm^64^. Uniquely mapping reads were filtered (MAPQ >10), PCR duplicates removed and mitochondrial reads discarded with SAMTools^65^. Downstream analysis was performed using BEDTools and deepTools^66^.

### Immunohistochemistry

Cortical organoids were harvested, washed twice with PBS, and fixed overnight in 4% paraformaldehyde (PFA) at 4°C. Following two additional PBS washes, organoids were cryoprotected in 30% sucrose overnight until they sank, then briefly washed with PBS before embedding in OCT compound (Fisher Scientific, 15212776) using an ethanol/dry ice slurry. Embedded organoids were stored at −80°C until sectioning. Cryosections of 20 μm were cut using a cryostat and collected on Superfrost slides, and stored at -80°C to ensure adhesion. For immunostaining, slides were brought to RT, sections were circled with a hydrophobic pen, and rehydrated with two 10-minute PBS washes. Sections were blocked and permeabilized in 1% BSA/0.3% Triton-X-100/PBS for 1 hour at RT in a humidity chamber, followed by overnight incubation with primary antibodies diluted in 1:3 blocking solution in PBS or PBS only for negative controls at 4°C. The next day, after three 10-minute PBS washes, sections were incubated with secondary antibodies (1:500) for 1 hour at RT, washed, and counterstained with DAPI. Slides were mounted with Prolong Diamond (Fisher Scientific, 15205739), and stored at 4°C until imaging. Antibodies used were PAX6 (Abcam ab5790, 1:150), FOXG1 (Takara M227, 1:150), SOX2 (Cambridge Bioscience 39843, 1:500), TUJ1 (Abcam MAB1195, 1:150), TBR1 (Abcam AB31940-1001, 1:200), CTIP2 (Abcam AB18465-100ug, 1:200).

### RT-qPCR

Total RNA was extracted using the Monarch Total RNA Miniprep Kit (New England BioLabs, T2010) according to the manufacturer’s instructions, including 15 min on-column DNase I treatment to remove genomic DNA. RNA was eluted in RNase-free ddH₂O and stored at −80°C until use. Reverse transcription was performed using the Maxima First Strand cDNA Synthesis Kit (25°C for 10 min, 50°C for 15 min, 85°C for 5 min). Gene expression was quantified by RT-qPCR using PowerUp SYBR Green Supermix (Life Technologies, A46112) on a CFX Connect Real-Time PCR Detection System (Bio-Rad) with the following cycling parameters: 95°C for 5 min, followed by 40 cycles of 95°C for 15 s, 60°C for 30 s, and 72°C for 30 s. GAPDH was used as a housekeeping gene for normalization. Primers are listed in the table below.

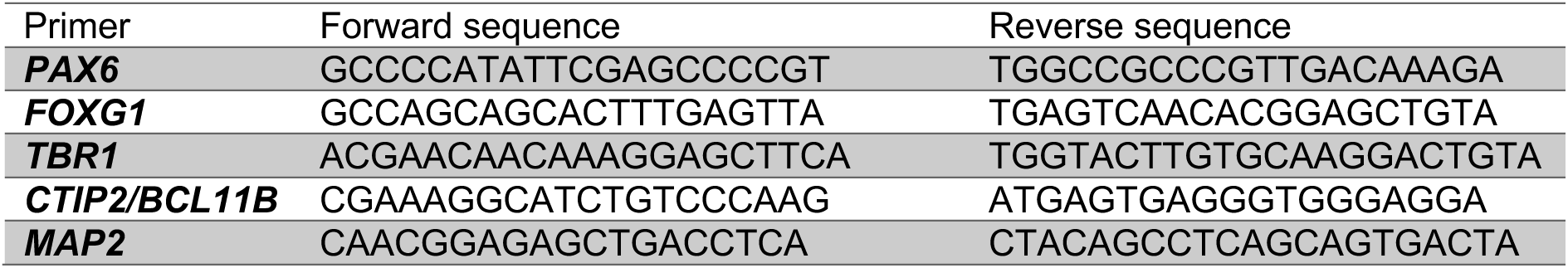
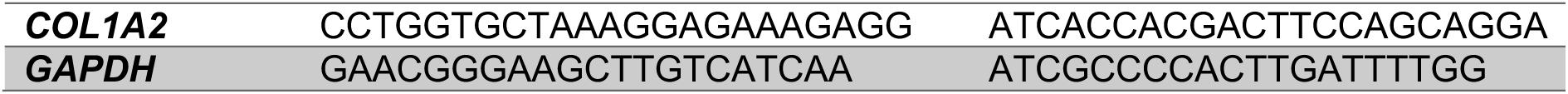

### Statistical analysis

All statistical analyses were performed in R (v4.5.1) or Python (v3.9.16). Specific statistical tests used for each analysis are described in the relevant sections above. Adjusted p-values were calculated using the Benjamini-Hochberg method unless otherwise stated. All experiments were performed with at least two independent biological replicates unless otherwise stated.

## Data availability

Raw and processed sequencing data is deposited in GEO under accession number GSE353922 (https://www.ncbi.nlm.nih.gov/geo/query/acc.cgi?acc=GSE353922).

## Acknowledgements

The authors are grateful to the Wysocka laboratory at Stanford University, particularly Dr. Raquel Fueyo, for generously providing the dCas9-KRAB vector. We also thank Andrew Isopi and Dr. Samantha Barnada (Thomas Jefferson University) for their contribution to the design of the sgRNA-containing plasmid. The authors further acknowledge the Facility for Imaging by Light Microscopy (FILM) at Imperial College London for providing access to imaging facilities and valuable technical support. The FILM facility is supported in part by the Wellcome Trust (grant 104931/Z/14/Z) and the Biotechnology and Biological Sciences Research Council (BBSRC; grant BB/L015129/1). We would additionally like to acknowledge the NIHR Imperial BRC Genomics Facility for the processing of our 10X Genomics samples. This project was supported by funding from M.T. by the BBSRC (grant BB/Y000854/1).

## Author contributions

L.D: Conceptualization, Investigation, Formal analysis, Visualization, Data curation, Methodology, Writing – original draft. S.M: Investigation, Data curation, Writing – original draft. P.U: Methodology, Data curation. T.T.R: Investigation, Data curation. Z.H.M: Investigation, Writing - review & editing. M.D: Investigation, Writing - review & editing. A.K: Investigation, Writing - review & editing. X.L: Investigation, Writing - review & editing. K.B: Methodology, Writing - review & editing. R.B: Methodology, Resources, Writing – review & editing. M.T: Conceptualization, Funding acquisition, Supervision, Project administration, Investigation, Writing – review & editing. All authors read and approved the final manuscript.

---

**Supplementary Figure 1.**
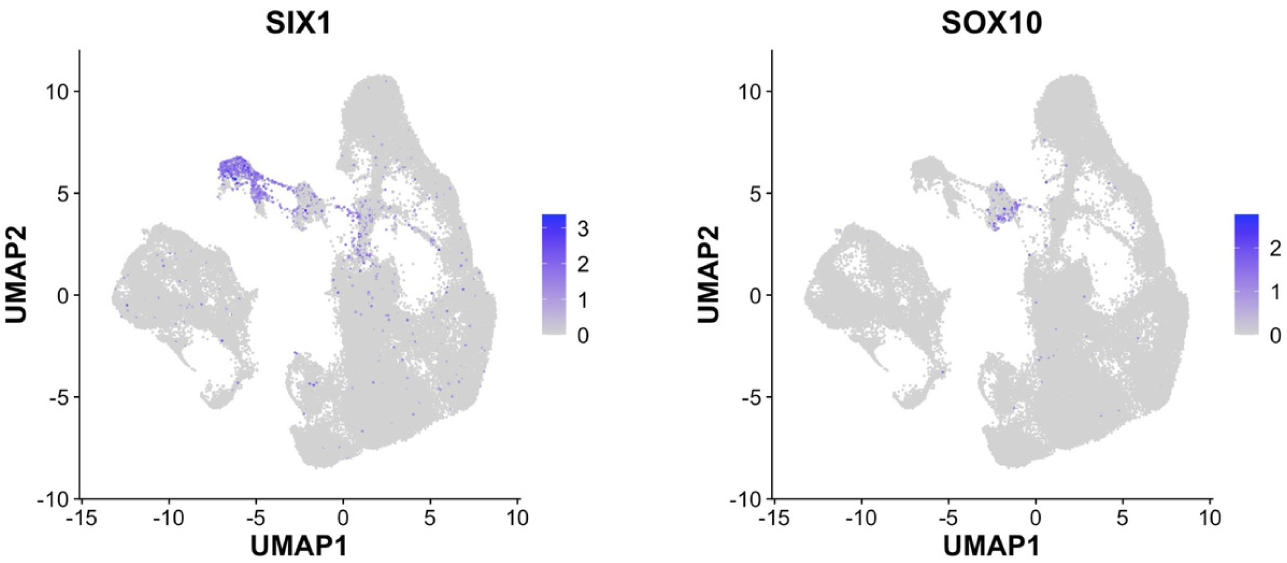
Marker gene expression of off-target cell populations in d21 cortical organoids. UMAP feature plots showing the expression of off-target population markers SIX1 and SOX10 in the short-read scRNA-seq dataset, with expression level indicated by colour intensity (white to dark blue). SIX1 marks placode-derived cells and SOX10 marks neural crest-derived cells; both populations were identified as minor off-target clusters and excluded from all downstream analyses

**Supplementary Figure 2.**
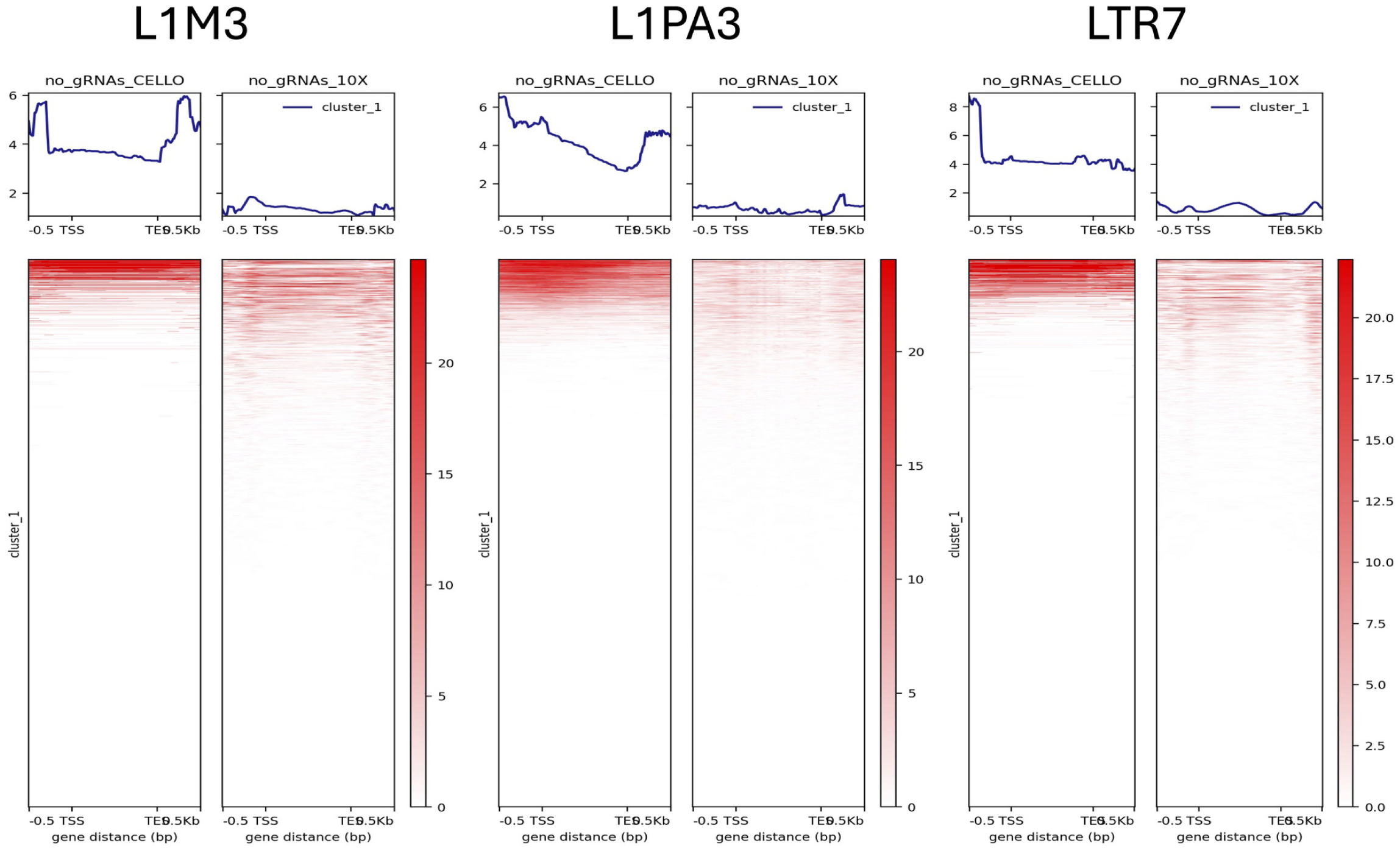
CELLO-seq improves detection of older TE subfamilies compared to short-read sequencing. Heatmaps of read coverage for three additional TE subfamilies: L1M3 (left), L1PA3 (middle), and LTR7 (right), com aring CELLO-seq and short-read scRNA-seq signal, aligned from 0.5 kb upstream of the TSS to TE end plus 0.5 kb downstream. Aggregate coverage profiles are shown above each heatmap. For all three subfamilies, CELLO-seq shows substantially higher read coverage of each element compared to short-read sequencing. These older TE subfamilies show an average four-fold increase in detection signal with CELLO-seq, demonstrating that the improvement in TE detection is not restricted to the youngest human-specific elements but extends across evolutionarily older subfamilies.

**Supplementary Figure 3.**
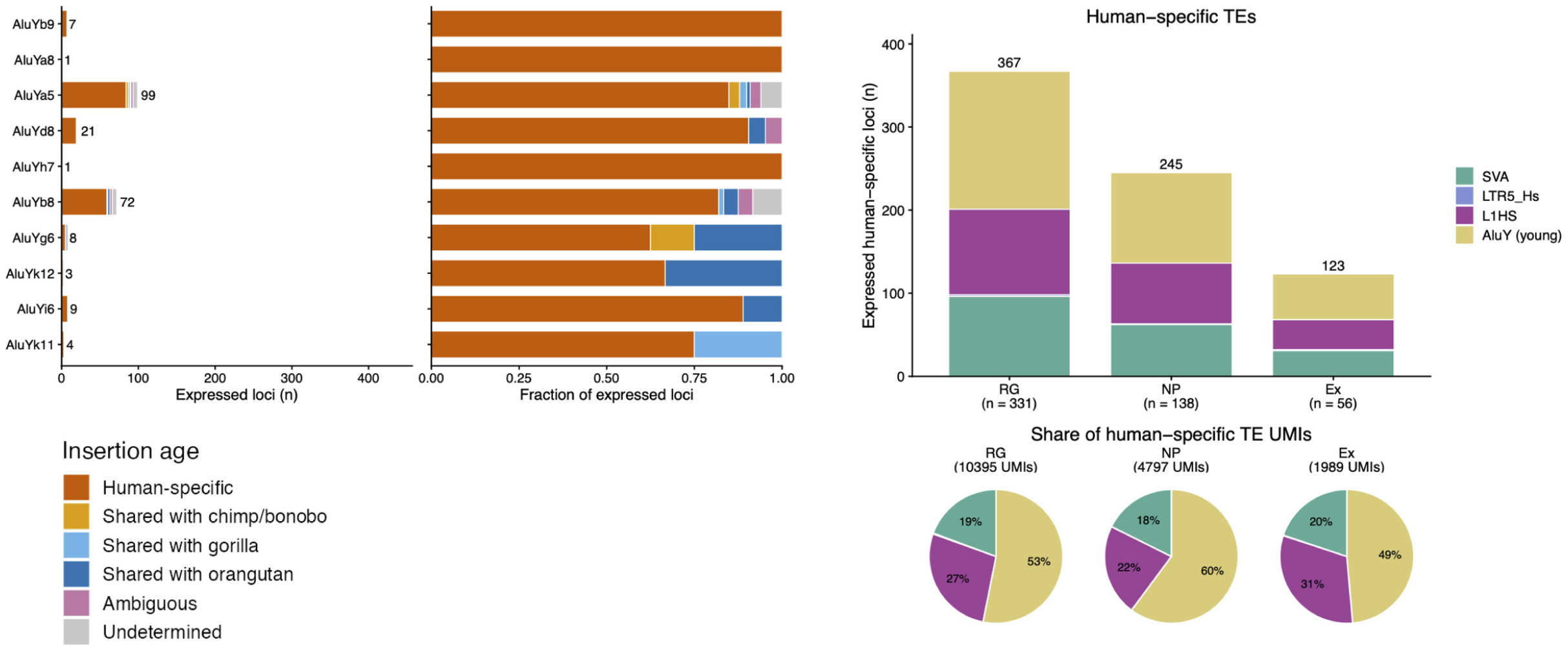
Expression of young AluY subfamily members in cortical organoids. Left: Bar chart showing the number of expressed loci per AluY subfamily member (AluYb9, AluYaδ, AluYaδ, AluYdδ, AluYh7, AluYbδ, AluYgδ, AluYk12, AluYiδ, AluYk11), with bars coloured by insertion age: human-specific (dark orange), shared with chimpanzee/bonobo (light orange), shared with gorilla (light blue), shared with orangutan (dark blue), ambiguous (pink), and undetermined (grey). The majority of expressed AluY loci carry human-specific insertions, with AluYaδ (n=99) and AluYbδ (n=72) being the most abundantly detected subfamilies. Middle: Stacked bar chart showing the proportional contribution of each insertion age category to expressed loci within each AluY subfamily, demonstrating the predominantly human-specific origin of detected elements. Right: Bar chart (top) showing the total number of expressed human-specific TE loci per cell type including AluY elements alongside SVA (teal), LTRδHs (green), L1 Hs (purple), and AluY young (yellow) families, with RG expressing the most loci (367 total), followed by NP (245) and Ex (123). Pie charts (bottom) showing the proportional UMI contribution of each TE family per cell type (RG: 10,395 UMIs; NP: 4,797 UMIs; Ex: 1,9δ9 UMIs), revealing that when AluY elements are included, they account for a dominant fraction of detected TE UMIs across all cell types. As Alu elements are transcribed by RNA Polymerase III and lack poly-A tails, their detection by poly-A capture-based CELLO-seq likely reflects read-through transcription from upstream Pol II loci and they were therefore excluded from the main analyses

**Supplementary Figure 4.**
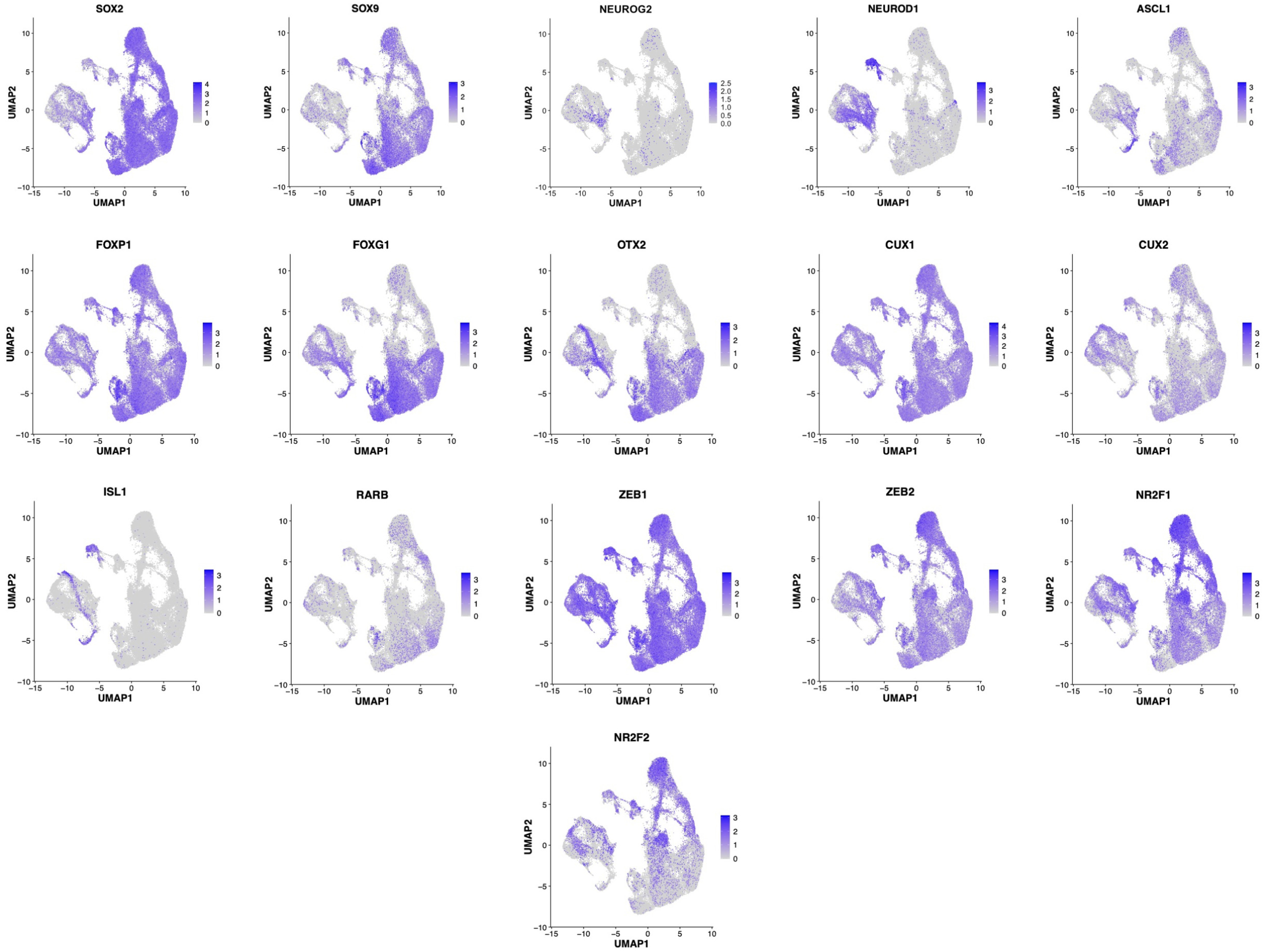
Distribution of TF expression following motif enrichment analysis in cortical d21 organoids. UMAP visualisations of the expression levels of transcription factors whose motifs were found enriched in cortical organoids. These motifs were enriched in either radial glia, neural progenitors, excitatory neurons or a combination. Expressing cells were ordered in front in order to visualise lowly expressed genes.

**Supplementary Figure 5.**
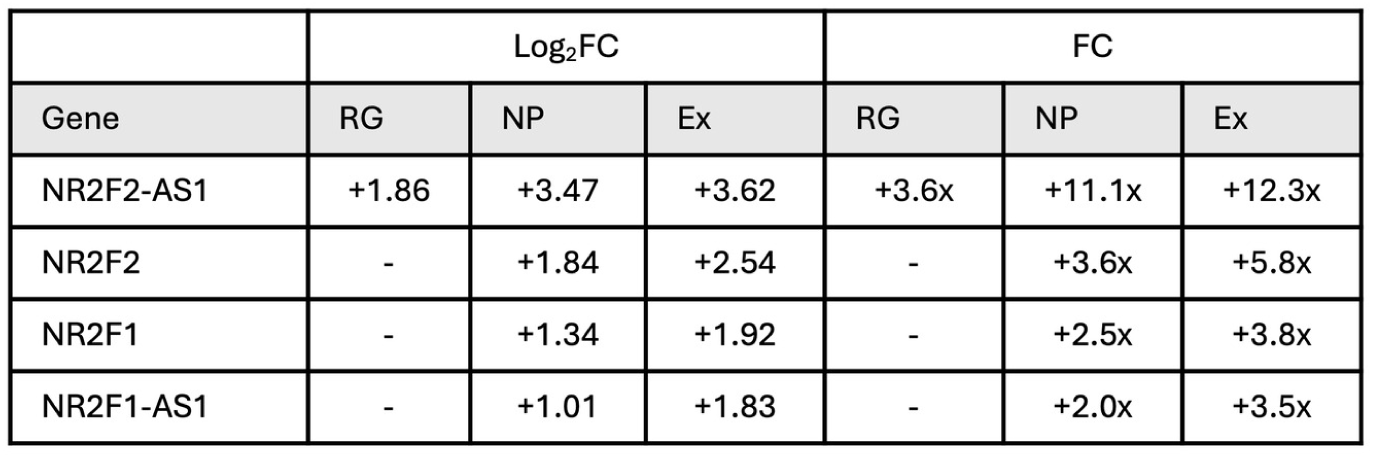
Progressive gene upregulation of NR2F1 and NR2F2 and their antisense transcripts along the differentiation axis. Average log2 fold changes (Log2FC) and fold changes (FC) in gene expression shown between CRISPRI (÷gRNAs) and control cells (-gRNAs) at day 21 of cortical organoid differentiation (n=2). All fold changes shown were statistically significant (FDR < 0.05).

**Supplementary Figure 6.**
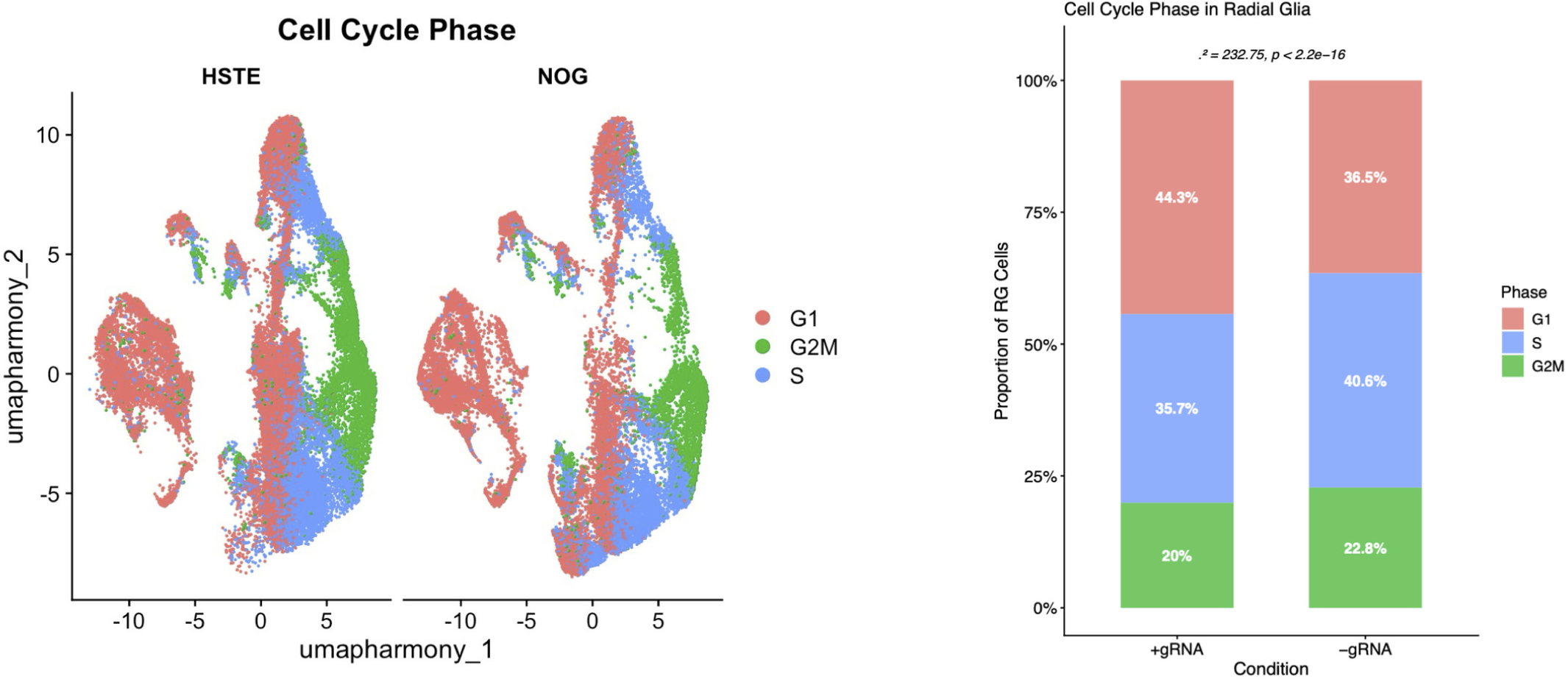
Cell cycle phase distribution in CRISPRi and control cortical organoids. Left: UMAP visualisation of cell cycle phase scoring for CRISPRi (÷gRNA, HSTE, left) and control (-gRNA, NOG, right) organoids, with cells coloured by inferred cell cycle phase: G1 (salmon), S (blue), and G2M (green). In both conditions, the radial glia cluster shows a spatially ordered distribution of cell cycle phases on the UMAP, consistent with active cell cycle progression and sequential transitions from S phase through G2M and back to G1. Neuronal populations are predominantly in G1 in both conditions, confirming the expected post-mitotic state of differentiated neurons and the overall directionality of the differentiation trajectory. Right: Stacked bar chart showing the proportion of radial glia cells in each cell cycle phase for CRISPRi (÷gRNA) and control (-gRNA) conditions.

## References

1. Trizzino, M., Kapusta, A. & Brown, C. D. Transposable elements generate regulatory novelty in a tissue-specific fashion. BMC Genomics 19, 468 (2018).

2. Fueyo, R., Judd, J., Feschotte, C. & Wysocka, J. Roles of transposable elements in the regulation of mammalian transcription. Nat. Rev. Mol. Cell Biol. 23, 481–497 (2022).

3. Lynch, V. J., Leclerc, R. D., May, G. & Wagner, G. P. Transposon-mediated rewiring of gene regulatory networks contributed to the evolution of pregnancy in mammals. Nat. Genet. 43, 1154–1159 (2011).

4. Bourque, G. et al. Evolution of the mammalian transcription factor binding repertoire via transposable elements. Genome Res. 18, 1752–1762 (2008).

5. Kunarso, G. et al. Transposable elements have rewired the core regulatory network of human embryonic stem cells. Nat. Genet. 42, 631–634 (2010).

6. Sundaram, V. & Wysocka, J. Transposable elements as a potent source of diverse cis-regulatory sequences in mammalian genomes. Philosophical Transactions of the Royal Society B: Biological Sciences 375, 20190347 (2020).

7. Schmidt, D. et al. Waves of retrotransposon expansion remodel genome organization and CTCF binding in multiple mammalian lineages. Cell 148, 335–348 (2012).

8. Jacques, P.-É., Jeyakani, J. & Bourque, G. The Majority of Primate-Specific Regulatory Sequences Are Derived from Transposable Elements. PLoS Genet. 9, e1003504 (2013).

9. Chuong, E. B., Rumi, M. A. K., Soares, M. J. & Baker, J. C. Endogenous retroviruses function as species-specific enhancer elements in the placenta. Nat. Genet. 45, 325– 329 (2013).

10. Chuong, E. B., Elde, N. C. & Feschotte, C. Regulatory evolution of innate immunity through co-option of endogenous retroviruses. Science (1979). 351, 1083–1087 (2016).

11. Pontis, J. et al. Primate-specific transposable elements shape transcriptional networks during human development. Nat. Commun. 13, 1–13 (2022).

12. Patoori, S., Barnada, S. M., Large, C., Murray, J. I. & Trizzino, M. Young transposable elements rewired gene regulatory networks in human and chimpanzee hippocampal intermediate progenitors. Development (Cambridge) 49, (2022).

13. Barnada, S. M. et al. Genomic features underlie the co-option of SVA transposons as cis-regulatory elements in human pluripotent stem cells. PLoS Genet. 18, 1–19 (2022).

14. Deelen, L. et al. Hominoid-specific transposable elements reshaped neural crest migration in craniofacial development. Mol. Syst. Biol. 21, 1731–1747 (2025).

15. Cordaux, R. & Batzer, M. A. The impact of retrotransposons on human genome evolution. Nat. Rev. Genet. 10, 691–703 (2009).

16. Iskow, R. C. et al. Natural mutagenesis of human genomes by endogenous retrotransposons. Cell 141, 1253–1261 (2010).

17. Konkel, M. K. et al. Sequence analysis and characterization of active human alu subfamilies based on the 1000 genomes pilot project. Genome Biol. Evol. 7, 2608– 2622 (2015).

18. Beck, C. R. et al. LINE-1 retrotransposition activity in human genomes. Cell 141, 1159–1170 (2010).

19. Subramanian, R. P., Wildschutte, J. H., Russo, C. & Coffin, J. M. Identification, characterization, and comparative genomic distribution of the HERV-K (HML-2) group of human endogenous retroviruses. Retrovirology 8, 90 (2011).

20. Pontis, J. et al. Hominoid-Specific Transposable Elements and KZFPs Facilitate Human Embryonic Genome Activation and Control Transcription in Naive Human ESCs. Cell Stem Cell 24, 724–735.e5 (2019).

21. Lanciano, S. & Cristofari, G. Measuring and interpreting transposable element expression. Nature Reviews Genetics vol. 21 721–736 Preprint at 10.1038/s41576-020-0251-y (2020).

22. Berrens, R. V. et al. Locus-specific expression of transposable elements in single cells with CELLO-seq. Nat. Biotechnol. 40, 546–554 (2022).

23. Marlow, S. A., Deaville, L. A. & Berrens, R. V. Long-read RNA sequencing of transposable elements from single cells using CELLO-seq. Nat. Protoc. 20, 3070– 3095 (2025).

24. Vanderhaeghen, P. & Polleux, F. Developmental mechanisms underlying the evolution of human cortical circuits. Nat. Rev. Neurosci. 24, 213–232 (2023).

25. Benito-Kwiecinski, S. et al. An early cell shape transition drives evolutionary expansion of the human forebrain. Cell 184, 2084–2102.e19 (2021).

26. Playfoot, C. J. et al. Transposable elements and their KZFP controllers are drivers of transcriptional innovation in the developing human brain. Genome Res. 31, 1531– 1545 (2021).

27. Bertucci, T. et al. Integrated manufacturing platform for cortical organoids reveals early phosphatase and prefoldin dysregulation in MAPT V337M neurons. npj Dementia 2, 100 (2026).

28. Polioudakis, D. et al. A Single-Cell Transcriptomic Atlas of Human Neocortical Development during Mid-gestation. Neuron 103, 785–801.e8 (2019).

29. Bowles, K. R. et al. ELAVL4, splicing, and glutamatergic dysfunction precede neuron loss in MAPT mutation cerebral organoids. Cell 184, 4547–4563.e17 (2021).

30. Eze, U. C., Bhaduri, A., Haeussler, M., Nowakowski, T. J. & Kriegstein, A. R. Single-cell atlas of early human brain development highlights heterogeneity of human neuroepithelial cells and early radial glia. Nat. Neurosci. 24, 584–594 (2021).

31. Wang, H. et al. SVA elements: A hominid-specific retroposon family. J. Mol. Biol. 354, 994–1007 (2005).

32. Padmanabhan Nair, V., et al. Activation of HERV-K(HML-2) disrupts cortical patterning and neuronal differentiation by increasing NTRK3. Cell Stem Cell 28, 1566–1581.e8 (2021).

33. Adami, A. et al. LINE-1 retrotransposons mediate cis-acting transcriptional control in human pluripotent stem cells and regulate early brain development. Cell Genomics 5, (2025).

34. Toda, T. et al. Long interspersed nuclear elements safeguard neural progenitors from precocious differentiation. Cell Rep. 43, (2024).

35. Garza, R. et al. LINE-1 retrotransposons drive human neuronal transcriptome complexity and functional diversification. Sci. Adv. 9, eadh9543 (2023).

36. Naka, H., Nakamura, S., Shimazaki, T. & Okano, H. Requirement for COUP-TFI and II in the temporal specification of neural stem cells in CNS development. Nat. Neurosci. 11, 1014–1023 (2008).

37. Lui, J. H., Hansen, D. V. & Kriegstein, A. R. Development and evolution of the human neocortex. Cell vol. 146 18–36 Preprint at 10.1016/j.cell.2011.06.030 (2011).

38. Lindhout, F. W., Krienen, F. M., Pollard, K. S. & Lancaster, M. A. A molecular and cellular perspective on human brain evolution and tempo. Nature 630, 596–608 (2024).

39. Florio, M. & Huttner, W. B. Neural progenitors, neurogenesis and the evolution of the neocortex. Development 141, 2182–2194 (2014).

40. Sultan, K. T., Brown, K. N. & Shi, S. H. Production and organization of neocortical interneurons. Frontiers in Cellular Neuroscience vol. 7 Preprint at 10.3389/fncel.2013.00221 (2013).

41. Radonjić, N. V. et al. Diversity of Cortical Interneurons in Primates: The Role of the Dorsal Proliferative Niche. Cell Rep. 9, 2139–2151 (2014).

42. Yu, X. & Zecevic, N. Dorsal radial glial cells have the potential to generate cortical interneurons in human but not in mouse brain. Journal of Neuroscience 31, 2413– 2420 (2011).

43. Letinic, K., Zoncu, R. & Rakic, P. Origin of GABAergic neurons in the human neocortex. Nature 417, 645–649 (2002).

44. Yokota, Y. et al. Radial glial dependent and independent dynamics of interneuronal migration in the developing cerebral cortex. PLoS One 2, (2007).

45. Stewart, C. et al. A comprehensive map of mobile element insertion polymorphisms in humans. PLoS Genet. 7, (2011).

46. Witherspoon, D. J. et al. Mobile element scanning (ME-Scan) identifies thousands of novel Alu insertions in diverse human populations. Genome Res. 23, 1170–1181 (2013).

47. Gardner, E. J. et al. The mobile element locator tool (MELT): Population-scale mobile element discovery and biology. Genome Res. 27, 1916–1929 (2017).

48. Goubert, C., Zevallos, N. A. & Feschotte, C. Contribution of unfixed transposable element insertions to human regulatory variation. Philosophical Transactions of the Royal Society B: Biological Sciences 375, 20190331 (2020).

49. van Bree, E. J. et al. A hidden layer of structural variation in transposable elements reveals potential genetic modifiers in human disease-risk loci. Genome Res. 32, 656– 670 (2022).

50. Jönsson, M. E., Garza, R., Johansson, P. A. & Jakobsson, J. Transposable Elements: A Common Feature of Neurodevelopmental and Neurodegenerative Disorders. Trends in Genetics vol. 36 610–623 Preprint at 10.1016/j.tig.2020.05.004 (2020).

51. Wang, L., Norris, E. T. & Jordan, I. K. Human retrotransposon insertion polymorphisms are associated with health and disease via gene regulatory phenotypes. Front. Microbiol. 8, (2017).

52. Payer, L. M. et al. Structural variants caused by Alu insertions are associated with risks for many human diseases. Proc. Natl. Acad. Sci. U. S. A. 114, E3984–E3992 (2017).

53. Nadler, M. J. S. et al. Hominoid SVA-lncRNA AK057321 targets human-specific SVA retrotransposons in SCN8A and CDK5RAP2 to initiate neuronal maturation. *Commun*. Biol. 6, (2023).

54. Horváth, V. et al. Mini-heterochromatin domains constrain the cis-regulatory impact of SVA transposons in human brain development and disease. Nat. Struct. Mol. Biol. 31, 1543–1556 (2024).

55. Dias, C. et al. BCL11A Haploinsufficiency Causes an Intellectual Disability Syndrome and Dysregulates Transcription. Am. J. Hum. Genet. 99, 253–274 (2016).

56. Lessel, D. et al. BCL11B mutations in patients affected by a neurodevelopmental disorder with reduced type 2 innate lymphoid cells. Brain 141, 2299–2311 (2018).

57. Südhof, T. C. Neuroligins and neurexins link synaptic function to cognitive disease. Nature vol. 455 903–911 Preprint at 10.1038/nature07456 (2008).

58. Alarcón, M. et al. Linkage, Association, and Gene-Expression Analyses Identify CNTNAP2 as an Autism-Susceptibility Gene. Am. J. Hum. Genet. 82, 150–159 (2008).

59. Hong, S. E. et al. Autosomal recessive lissencephaly with cerebellar hypoplasia is associated with human RELN mutations. Nat. Genet. 26, 93–96 (2000).

60. Cao, J. et al. The single-cell transcriptional landscape of mammalian organogenesis. Nature 566, 496–502 (2019).

61. Liao, Y., Wang, J., Jaehnig, E. J., Shi, Z. & Zhang, B. WebGestalt 2019: gene set analysis toolkit with revamped UIs and APIs. Nucleic Acids Res. 47, W199–W205 (2019).

62. Morabito, S., Reese, F., Rahimzadeh, N., Miyoshi, E. & Swarup, V. hdWGCNA identifies co-expression networks in high-dimensional transcriptomics data. Cell Reports Methods 3, (2023).

63. Heinz, S. et al. Simple Combinations of Lineage-Determining Transcription Factors Prime cis-Regulatory Elements Required for Macrophage and B Cell Identities. Mol. Cell 38, 576–589 (2010).

64. Li, H. & Durbin, R. Fast and accurate short read alignment with Burrows-Wheeler transform. Bioinformatics 25, 1754–1760 (2009).

65. Li, H. et al. The Sequence Alignment/Map format and SAMtools. Bioinformatics 25, 2078–2079 (2009).

66. Ramírez, F., Dündar, F., Diehl, S., Grüning, B. A. & Manke, T. deepTools: a flexible platform for exploring deep-sequencing data. Nucleic Acids Res. 42, W187–W191 (2014).

